# Cultivated *Kapaibacteriales* reveal contrasting lake and groundwater lifestyles built on shared metabolic dependencies

**DOI:** 10.64898/2026.11.05.764759

**Authors:** Hongjae Park, Sumin Kim, Jaeho Jang, Hyeonuk Sa, Miri S. Park, Ahyoung Choi, Ilnam Kang, Jang-Cheon Cho

## Abstract

Microbial lineages occupying contrasting habitats offer an opportunity to examine how habitat-associated genomic diversification coexists with evolutionarily conserved metabolic traits. Here, we investigated this relationship in *Kapaibacteriales*, a globally distributed but poorly characterized bacterial group. We obtained 29 axenic isolates representing three families and nine novel species from lakes and groundwater, providing the first cultivated members of the order from non-thermal habitats. Global metagenomic profiling and comparative genomics revealed marked differences in ecological distributions and genomic architectures between lake and groundwater populations. Lake populations were widespread, recurrently enriched during summer, and had compact genomes. In contrast, groundwater populations were geographically restricted and had larger genomes enriched in genes for signal transduction, transcriptional regulation, and mobile genetic elements. Despite this divergence, both groups shared predicted deficiencies in the biosynthesis of multiple amino acids, vitamins, and respiratory cofactors, including quinones and heme. These findings indicate that greater coding capacity need not confer greater biosynthetic self-sufficiency, and that metabolic dependencies can persist through ecological diversification, with implications for cultivating abundant but uncultured aquatic lineages.

## Introduction

Microbial genomes are shaped by both ecological selection and evolutionary inheritance. Across aquatic microorganisms, variation in genome size and gene content has been associated with trophic strategies and environmental gradients, illustrating how genomic repertoires can reflect the conditions under which populations persist and grow^1–3^. Gene content, in turn, changes with genome size in a predictable manner, with regulatory and signal transduction genes increasing disproportionately in larger genomes^4,5^, and auxotrophies, which are widespread among bacteria^6^, being particularly extensive in smaller genomes^7^. Resource availability can also alter the costs and benefits of maintaining biosynthetic functions. Experimental studies have shown that loss of biosynthetic genes can confer a fitness advantage when the corresponding metabolites are consistently available from external sources^6^. More broadly, the Black Queen Hypothesis (BQH) provides a framework for understanding how metabolic dependencies evolve when costly functions performed by some community members generate public goods that others can exploit^8^. Yet genomic traits are not determined by ecology alone, as genome size and functional repertoires are often strongly structured by shared ancestry across prokaryotes^9^. A central challenge in microbial comparative genomics is therefore to distinguish genomic features that vary with habitat from those that remain constrained by evolutionary history.

Comparisons within a single lineage that spans contrasting habitats provide an effective way to address this challenge. Lake surface waters and groundwater represent such a contrast. Lake epilimnion is illuminated, seasonally variable in temperature, hydrologically connected, and supplied with phytoplankton-derived organic matter in recurrent pulses^10^, on which many heterotrophic bacteria, including abundant auxotrophic lineages, depend^11,12^. Aquifers, in contrast, are dark, thermally stable and hydrologically isolated, with spatially heterogeneous and often episodic resource inputs, and their microbial communities are frequently structured by dispersal limitation and strong site specificity^10,13^. Although groundwater harbors lineages with extremely reduced genomes, such as *Patescibacteriota^14^*, larger genomes have also been reported in deeper aquifers^15,16^, leaving it unresolved how such conditions shape genome architecture. Closely related populations distributed across lakes and aquifers therefore provided an opportunity to test whether ecological differentiation is accompanied by changes in genome size, resource-acquisition capacity, regulatory complexity, and biosynthetic potential, or whether some metabolic traits remain conserved despite environmental divergence.

Resolving these alternatives requires more than comparisons of metagenome-derived genomes alone. Environmental surveys are needed to establish where populations occur and under which conditions, whereas cultivated representatives provide genomic and organismal resolution that environmental genomes cannot. Metagenome-assembled genomes (MAGs) and single-amplified genomes (SAGs) are often incomplete or fragmented, which constrains inferences that depend on gene absence or copy number, such as predicted auxotrophies, paralog expansion, and prophage content^17,18^. Near-complete genomes from axenic isolates allow such features to be resolved with greater confidence, while the cultures themselves enable physiological and morphological characterization. In abundant aquatic bacteria, cultivation has linked genomic deficiencies to specific growth requirements, such as the requirement of *Pelagibacter* (SAR11 clade) for exogenous reduced sulfur^19^, the reliance of *Prochlorococcus* on co-occurring microorganisms for hydrogen peroxide removal^20^, and the heme auxotrophy of the freshwater actinobacterial lineage acI (*Nanopelagicales*), whose growth in culture depends on supplementation with catalase, a heme-containing enzyme^21,22^. Integrating environmental surveys, isolate genomes, and physiology can therefore reveal which components of microbial genomes are ecologically flexible and which remain conserved across habitats.

The order *Kapaibacteriales* (GTDB R232; previously referred to as the candidate phylum “*Ca.* Kapaibacteriota”) is a globally distributed but poorly characterized bacterial lineage that is well suited to this approach^23^. First identified in 1998 as the 16S rRNA gene clone group OPB56 in a Yellowstone hot spring^24^, this lineage has since been detected from diverse aquatic, terrestrial, and engineered systems^25–27^, and can reach high relative abundances in hot springs and engineered systems. Originally recognized as a deeply branching lineage related to the phylum *Chlorobi*, it has been characterized mainly through genome-resolved metagenomic studies, which suggested aerobic heterotrophic potential in some members^23^. Its only cultivated representative, however, is an anaerobic thermophile recently isolated from deep thermal waters^28^, leaving aerobic and mesophilic freshwater members that are widespread in environmental surveys without cultured representatives. Consequently, how genome architecture and functional repertoires vary across habitats within this lineage, and whether genome-based predictions hold in living cells, remain unknown. Cultivating additional representatives from contrasting environments would provide a foundation for interpreting genomic variation in the context of ecological distributions and observable phenotypic traits.

In this study, we established 29 axenic isolates of *Kapaibacteriales* from lakes and groundwater, representing three families, four genera, and nine species, and used them to examine genomic, physiological, and morphological differentiation across the two contrasting habitats. We combined global metagenomic profiling with comparative analyses of isolate genomes and publicly available MAGs to determine how ecological distributions relate to genome size, regulatory complexity, and biosynthetic potential. We further characterized representative isolates physiologically and morphologically to examine whether ecological and genomic differentiation is accompanied by corresponding variation in growth and cellular phenotypes. We show that groundwater populations have larger genomes enriched in regulatory and mobile genetic elements, yet share with lake populations widespread predicted deficiencies in amino acid, vitamin, and respiratory cofactor biosynthesis.

## Results

### First cultivation of *Kapaibacteriales* from non-thermal environments

Using dilution-to-extinction cultivation in a low-nutrient freshwater medium (LNFM; Table S1), we obtained 29 axenic isolates of *Kapaibacteriales* (Table S2) from three lakes and one aquifer in the Republic of Korea (Fig. 1a and Table S3), namely an oligotrophic lake (Lake Soyang, *n* = 4), two eutrophic lakes (Lake Daechung, *n* = 13; Lake Paldang, *n* = 1), and a groundwater well (Yanggu, *n* = 11). Apart from a single thermophilic species, *Eximiradius proteiniborus*, described recently^28^, the order *Kapaibacteriales* previously lacked cultivated representatives, making these 29 isolates the first from non-thermal environments and a substantial expansion of its cultured diversity. Near-complete genomes were obtained for all 29 isolates, with >99% completeness and <1.5% contamination (Table S2). GTDB-based classification assigned the isolates to three families (*Kapaibacteriaceae*, UBA961, and NICIL-2; Fig. 1b) and four genera, and clustering at 95% average nucleotide identity (ANI) resolved nine species (Table S4). Six of these species corresponded to GTDB placeholder species previously represented only by MAGs, whereas the remaining three species had no genomic representatives in GTDB. One of the four genera corresponds to *Kapaibacterium*, which has been registered under the SeqCode with a MAG as its nomenclatural type^23^. Here, we propose names for nine novel species, including *Kapaibacterium aquifericola* sp. nov., three novel genera (*Bradylimnobacter* gen. nov., *Dulcisphaera* gen. nov., and *Phreatorhabdus* gen. nov.), and two novel families, *Bradylimnobacteraceae* fam. nov. and *Phreatorhabdaceae* fam. nov., corresponding to the GTDB placeholder families UBA961 and NICIL-2, respectively. Formal descriptions of the new taxa and emendations of the family *Kapaibacteriaceae* and the genus *Kapaibacterium*, together with etymologies, are provided in Supplementary Note.

**Fig. 1.**
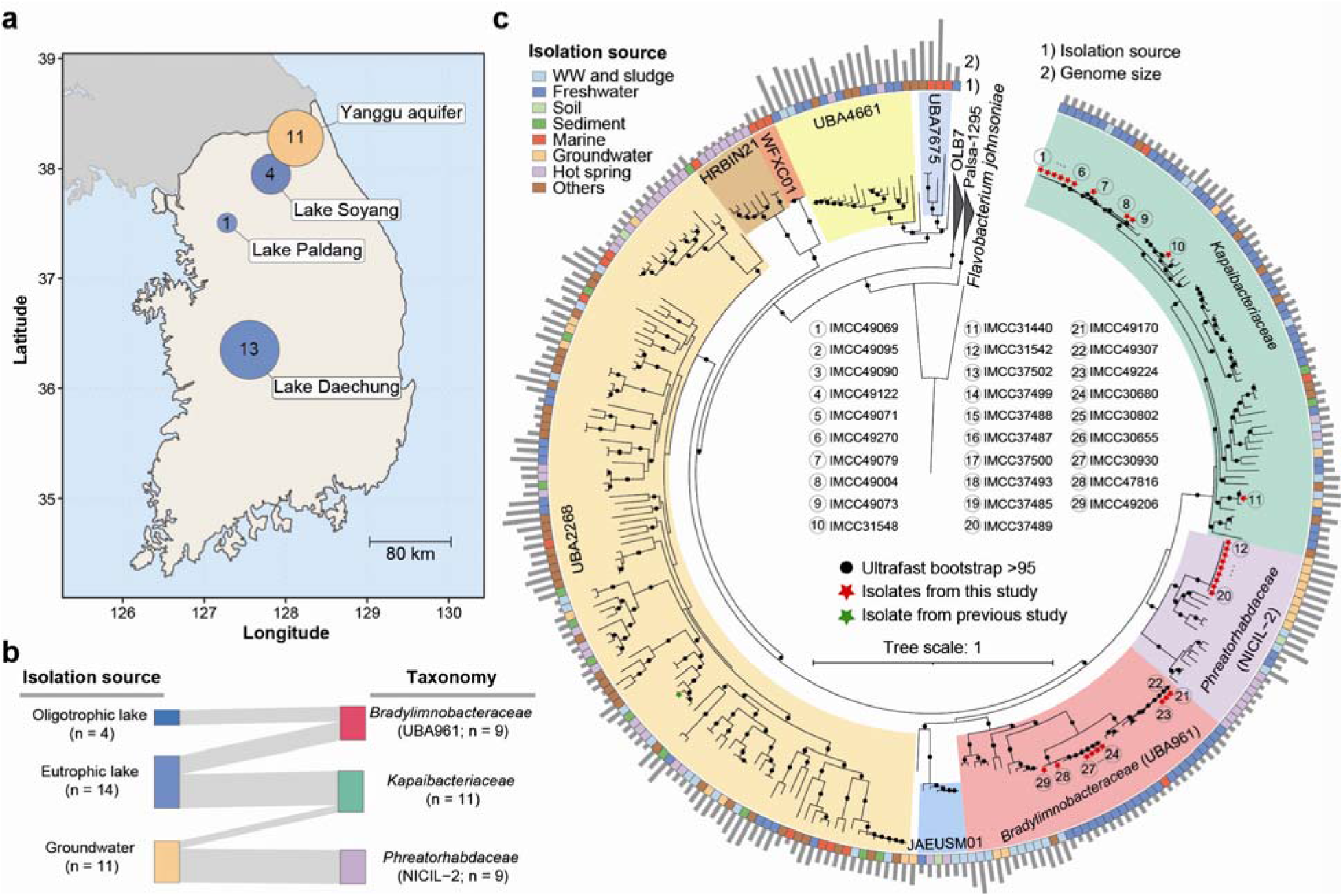
Genomic and geographic overview of *Kapaibacteriales* isolates. **(a)** Locations of the sampling sites in the Republic of Korea. Circle size and the number inside each circle indicate the number of isolate genomes obtained from each site. **(b)** Isolation source and family-level taxonomy of the 29 isolate genomes. **(c)** Maximum-likelihood phylogenomic tree of *Kapaibacteriales* isolate genomes and high-quality MAGs, constructed from a concatenated alignment of 120 bacterial single-copy marker proteins. Clades are shaded by family, and families proposed in this study are labeled with their GTDB placeholder names in parentheses. The inner ring indicates the isolation source and the outer bars indicate genome size. Black circles on nodes indicate ultrafast bootstrap support >95%. Red stars indicate isolates obtained in this study, numbered as listed in the center, and the green star indicates recently isolated *Eximiradius proteiniborus*. *Flavobacterium johnsoniae* was used as the outgroup. WW, wastewater.

To place the isolates within the broader genomic diversity of this lineage, we reconstructed a phylogenomic tree combining the 29 isolates, *E. proteiniborus*, and 248 high-quality MAGs (≥90% completeness and ≤5% contamination; Table S5) from GlobDB (Fig. 1c). These genomes originated from diverse habitat categories that were interspersed throughout the phylogeny, which may reflect repeated transitions between contrasting environments. The 30 isolate genomes span four heavily sampled families in the order, *Kapaibacteriaceae*, *Bradylimnobacteraceae*, *Phreatorhabdaceae*, and UBA2268, which together account for 83% (207/248) of the MAGs, providing cultured references for most of the known genomic diversity of the order. Notably, all previously available *Phreatorhabdaceae* MAGs originated from freshwater, hot springs, wastewater, or soil environments. The nine groundwater isolates, all belonging to *Phreatorhabdus flagellata*, are therefore the first *Phreatorhabdaceae* genomes recovered from an aquifer, expanding the known environmental breadth of this family and suggesting that additional diversity remains to be discovered in subsurface environments.

### Habitat-associated contrasts in genome size

Genome sizes of *Kapaibacteriales* differed systematically between lake and groundwater habitats (Fig. 2a, Table S2). *Kapaibacteriaceae* isolates were recovered from both habitats, enabling a direct within-family comparison. The two groundwater isolates, *K. aquifericola* IMCC31440 and *D. phreaticola* IMCC31548, had genomes of approximately 3.4 Mbp, up to 42% larger than those of the lake isolates (2.4–2.5 Mbp), despite similarly high GC contents (51– 57%). This difference was also evident within a single genus, as the genome of *D. phreaticola* was approximately 0.9 Mbp larger than those of the three lake-derived *Dulcisphaera* species. In contrast, *Bradylimnobacteraceae* isolates were recovered only from lakes and were characterized by small genomes (2.4–2.6 Mbp) and low GC contents (33–40%), whereas *Phreatorhabdaceae* isolates were recovered only from groundwater and had the largest genomes in the collection (∼4.1 Mbp). Publicly available MAGs provided complementary habitat comparisons for these two families. The two groundwater *Bradylimnobacteraceae* MAGs had substantially larger genomes (3.4 and 4.1 Mbp) than both the lake isolates and the 13 freshwater MAGs of this family (2.2–2.5 Mbp), whereas the two freshwater *Phreatorhabdaceae* MAGs had smaller genomes (2.4 and 2.5 Mbp) than the groundwater isolates (Fig. 2a and Table S5). Thus, all three families showed the same direction of lake–groundwater differentiation, with larger genomes associated with groundwater members, although the single groundwater MAG of *Kapaibacteriaceae* had a small genome (2.5 Mbp; Fig. 1c). A similar pattern was evident across the broader collection of *Kapaibacteriales* MAGs, which encompassed additional families and habitats (Fig. S1). Genomes from groundwater and biofilm environments were generally larger, with median sizes of 3.5–3.6 Mbp, than those from hot spring and freshwater environments (2.4– 2.6 Mbp).

**Fig. 2.**
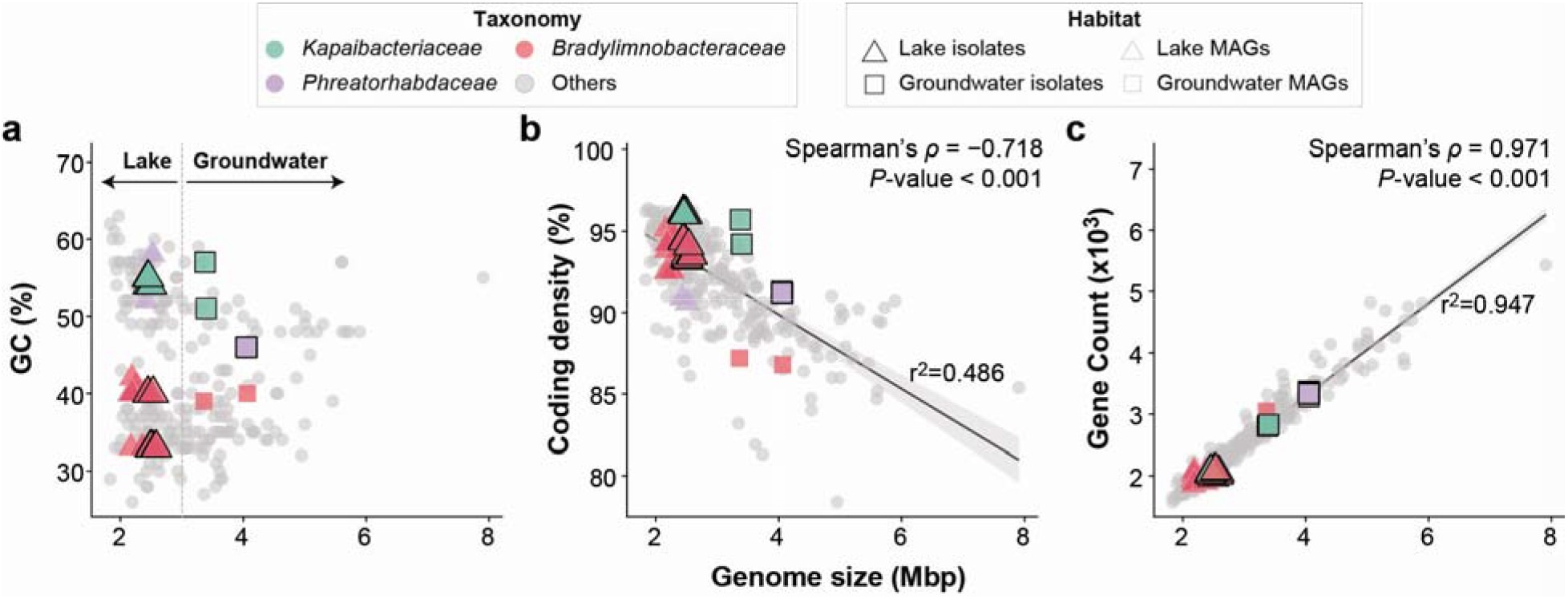
Genomic features of *Kapaibacteriales* isolates and MAGs in relation to genome size. Genome size is plotted against **(a)** GC content, **(b)** coding density and **(c)** gene count for the 29 isolates and 248 high-quality MAGs. Isolates (black outlines) are colored by family and shaped by habitat (triangles, lake; squares, groundwater). Lake and groundwater MAGs of *Bradylimnobacteraceae* and *Phreatorhabdaceae* are shown with the same colors and shapes without outlines for within-family comparison, while all other MAGs are shown as grey circles. In (a), the dashed line at 3.0 Mbp separates lake isolates (all <3.0 Mbp) from groundwater isolates (all >3.0 Mbp). In (b) and (c), Spearman’s rank correlation coefficients (_ρ_) and *P* values are shown. Solid lines represent linear regression fits, with 95% confidence intervals shaded in grey; r² denotes the coefficient of determination for each regression.

To examine how genome size variation was reflected in genome organization and gene content, we compared coding density and gene count across all isolate genomes and MAGs (Fig. 2b). Coding density decreased with increasing genome size (Spearman’s _ρ_ = −0.718, *P* < 0.001), although within-family habitat contrasts did not uniformly follow this relationship. In particular, freshwater and groundwater genomes of *Phreatorhabdaceae* had similar coding densities (∼91%) despite differing in genome size by approximately 1.5 Mbp. Gene count, in turn, was strongly correlated with genome size (Spearman’s _ρ_ = 0.971, *P* < 0.001; Fig. 2c), indicating that the larger groundwater genomes encode proportionally more genes rather than accumulating non-coding sequence.

### Physiological and morphological variation among isolates

To characterize the growth physiology and cellular morphology of *Kapaibacteriales*, we selected five representative isolates based on their ability to be reliably subcultured, their phylogenetic positions, and their ecological origins. All five isolates grew fastest at 25–30 °C, but their maximum growth rates and cell densities differed markedly (Figs. 3 and S2). The lake and groundwater isolates of *Kapaibacteriaceae* (IMCC49270 and IMCC31440) showed the highest growth rates (∼1.5 day^−1^) and reached maximum cell densities of 1.0–1.4 ×10^7^ cells mL^−1^. In contrast, the lake-derived *Bradylimnobacteraceae* isolate IMCC49307 grew most slowly (0.4 day^−1^) and reached the lowest maximum cell density (7.5 ×10^4^ cells mL^−1^), more than an order of magnitude lower than those of the other isolates, possibly reflecting an oligotrophic lifestyle or a requirement for growth factors not supplied in the medium. Notably, the two *Phreatorhabdaceae* isolates (IMCC31542 and IMCC37485) differed substantially in growth despite sharing >99.9% ANI. IMCC37485 reached a 50% higher maximum growth rate than IMCC31542 (0.9 vs. 0.6 day^−1^) and an order of magnitude higher maximum cell density (4.4 × 10^7^ vs. 2.6 × 10^6^ cells mL^−1^), indicating considerable physiological variation between genomically near-identical strains.

**Fig. 3.**
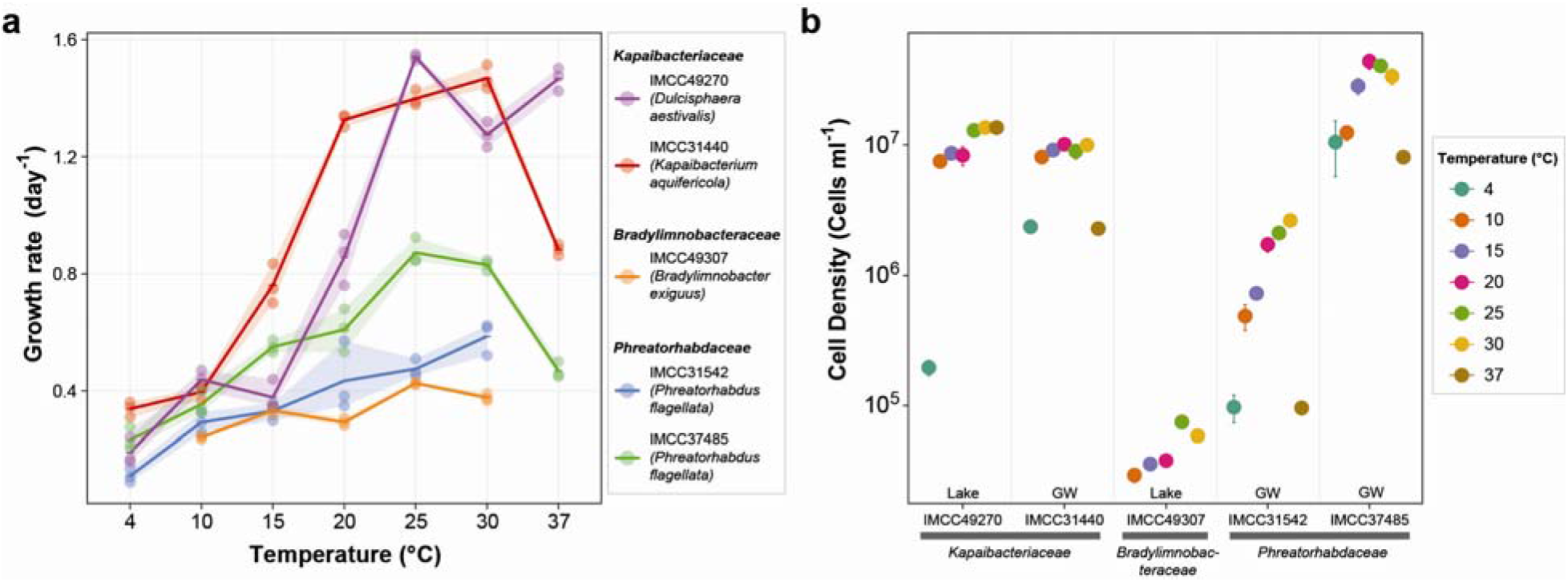
Temperature-dependent growth characteristics of five representative *Kapaibacteriales* isolates. **(a)** Specific growth rate (day ¹) of each isolate at temperatures from 4 to 37 °C. Points indicate individual cultures, and lines and shaded areas indicate the mean and range (minimum to maximum) of triplicate cultures. **(b)** Maximum cell density (cells mL ¹) of each isolate observed within 24 days at each temperature. Points and error bars indicate the mean and standard deviation of triplicate cultures. IMCC49307 was not tested at 4 or 37 °C. GW, groundwater. Time-course growth curves at each temperature are shown in Fig. S2.

The isolates also differed in their responses to temperatures outside their optima (Figs. 3 and S2). At 37 °C, the lake-derived *Kapaibacteriaceae* isolate IMCC49270 maintained a growth rate (1.5 day^−1^) and maximum cell density (1.4 × 10^7^ cells mL^−1^) comparable to those at its optimum, whereas its groundwater counterpart IMCC31440 was strongly inhibited, with its growth rate decreasing to 0.9 day^−1^ and its maximum cell density to 2.3 × 10^6^ cells mL^−1^. The two groundwater *Phreatorhabdaceae* isolates were likewise sensitive to elevated temperature. IMCC31542 showed no detectable growth at 37 °C, and IMCC37485 grew only after a lag of approximately 15 days, with its growth rate declining from 0.9 day^−1^ at 25 °C to 0.5 day^−1^ at 37 °C (Fig. S2). The opposite trend was observed at 4 °C, where the groundwater *Kapaibacteriaceae* isolate IMCC31440 reached 2.4 × 10^6^ cells mL ¹, whereas the lake isolate IMCC49270 showed little growth. These temperature responses were broadly consistent with the thermal regimes of the source environments, as lake surface temperatures reached up to 31.3 °C in summer, whereas groundwater temperatures remained at 13–14 °C throughout the sampling period (Table S3). Nevertheless, all groundwater isolates grew fastest at 25–30 °C, well above their *in situ* temperatures.

Scanning and transmission electron microscopy (SEM and TEM, respectively) revealed that cell morphology differed among the isolates in accordance with family affiliation (Fig. S3). *Kapaibacteriaceae* isolates were coccoid, 0.3–0.5 µm in diameter, whereas *Bradylimnobacteraceae* and *Phreatorhabdaceae* isolates were rod-shaped. The *Bradylimnobacteraceae* isolate IMCC49307 was particularly small (approximately 0.2 × 0.4 µm), while *Phreatorhabdaceae* cells were longer (approximately 0.2 × 2.0 µm). Flagella were observed in both *Phreatorhabdaceae* isolates but not in the *Kapaibacteriaceae* or *Bradylimnobacteraceae* isolates.

### Global distribution reveals niche partitioning in lakes and localization in groundwater

*Kapaibacteriales* were widely distributed across environments but differed markedly in abundance among habitats. Screening of 707,470 metagenomes in the Sandpiper database^29^ detected the order in 31,941 samples (4.5%), spanning at least 11 habitat categories across natural (aquatic and terrestrial), engineered (wastewater treatment, bioreactor, and aquaculture), and animal-associated environments (Fig. 4a and Table S6). Its distribution extended from hot springs to Arctic lakes, from marine to freshwater environments, and from wastewater to oligotrophic groundwater. Relative abundance was highest in hot springs (*n* = 545 samples; median 0.36% in samples where it was detected, maximum 18.9%), where *Kapaibacteriales* frequently ranked among the three most abundant prokaryotic orders (Fig. 4b and Table S6). In freshwater (*n* = 5,331; median 0.16%, maximum 8.3%), the most enriched samples were often associated with phytoplankton, including algal raceways and cyanobacterial cultures. In groundwater, by contrast, the order was detected in 650 samples, mostly at low relative abundances (median 0.03%, maximum 6.0%), and rarely ranked among the five most abundant orders.

**Fig. 4.**
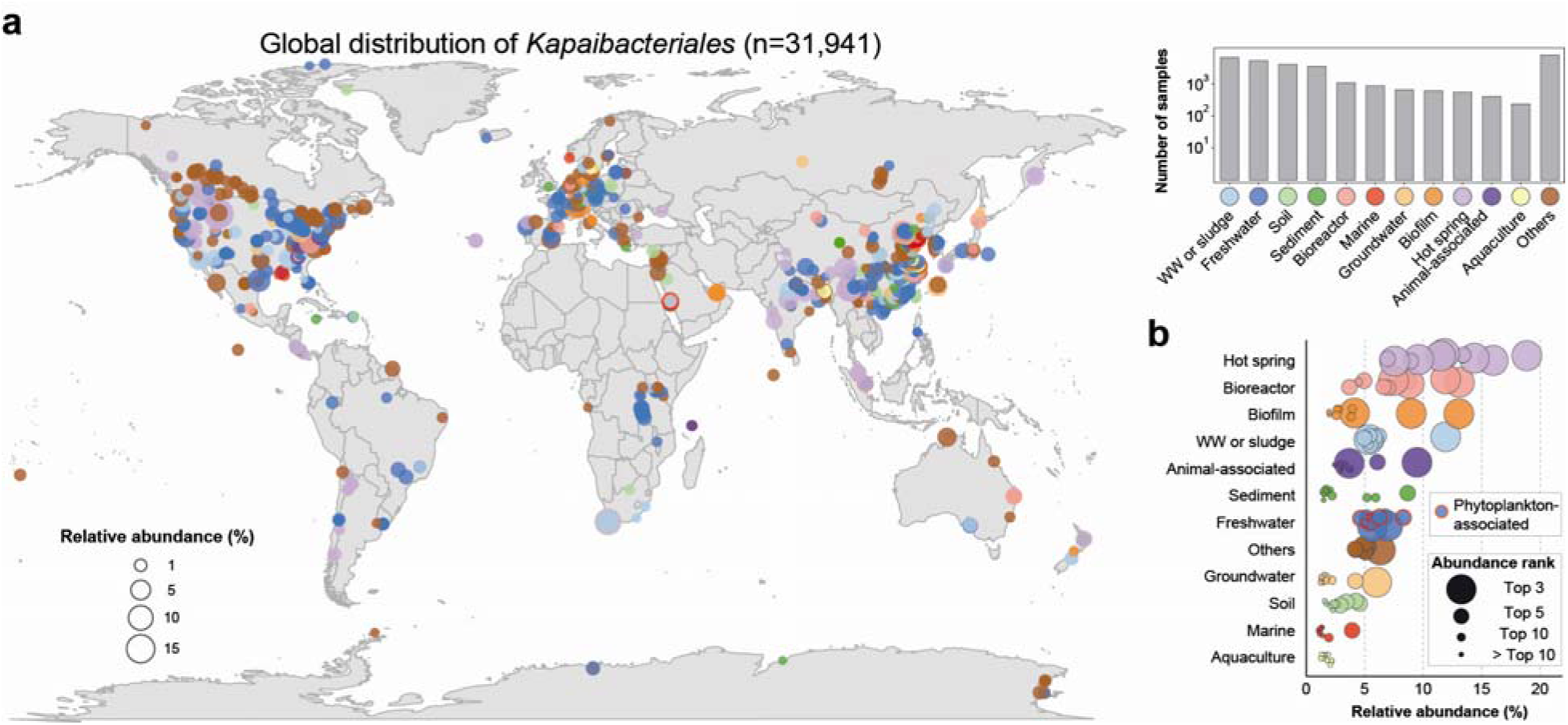
Global distribution and abundance of *Kapaibacteriales*. Data are from 31,941 metagenomes in the Sandpiper database in which *Kapaibacteriales* was detected. **(a)** Map of sampling sites, in which each circle represents a site with unique geographic coordinates. Where multiple metagenomes originated from the same site, the one with the highest relative abundance is shown. Circle size indicates the relative abundance (%) of *Kapaibacteriales*, and color indicates the habitat category. Only sites with ≥0.5% relative abundance are shown. The inset shows the number of metagenomes in each habitat category. **(b)** Relative abundance (%) of *Kapaibacteriales* in the 10 metagenomes with the highest *Kapaibacteriales* abundance in each habitat category. Circle size indicates the relative abundance rank of *Kapaibacteriales* among orders within each metagenome (top 3, top 5, top 10, or lower). Phytoplankton-associated metagenomes are highlighted with red outlines.

Within lakes, *Kapaibacteriales* showed pronounced seasonal dynamics. SingleM profiling^29^ of 2,067 freshwater metagenomes included multi-year time series from Lake Mendota (USA), Římov Reservoir (Czech Republic), Lake Biwa (Japan), and Trout Bog Lake (USA), representing eutrophic, meso-eutrophic, oligotrophic, and dystrophic conditions, respectively (Table S7). In these time series, *Bradylimnobacteraceae* and *Kapaibacteriaceae* dominated recurrent summer increases in the epilimnion (Fig. 5). In Lake Mendota, the relative abundance of *Kapaibacteriales* was positively correlated with that of cyanobacteria (Spearman’s _ρ_ = 0.555, *P* < 0.001), consistent with the coincident summer peaks of both groups observed across the 20-year time series. In contrast, neither family was detected in the dystrophic Trout Bog Lake or Crystal Bog Lake (Fig. 5 and Table S7), suggesting that their seasonal increases depend on conditions that are lacking in humic, dystrophic lakes.

**Fig. 5.**
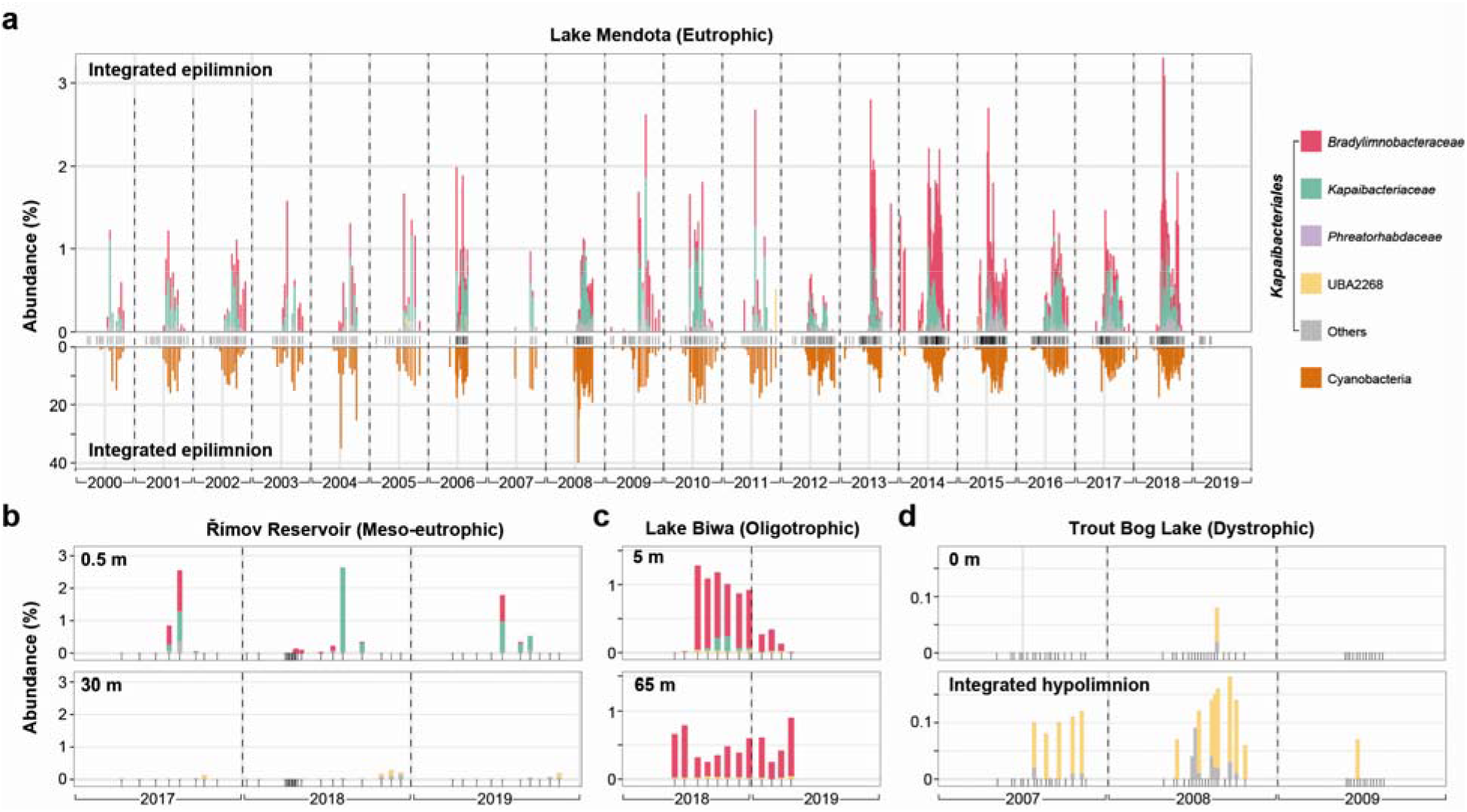
Seasonal distribution patterns of *Kapaibacteriales* in global lakes of varying trophic status. Relative abundance (%) of *Kapaibacteriales* families over time in **(a)** Lake Mendota (eutrophic; 2000–2019), **(b)** Římov Reservoir (meso-eutrophic; 2017–2019), **(c)** Lake Biwa (oligotrophic; 2018–2019), and (d) Trout Bog Lake (dystrophic; 2007–2009). Tick marks along the x-axis indicate sampling dates and dashed vertical lines separate years. For Lake Mendota, only integrated epilimnion samples were available, and the relative abundance of cyanobacteria in the same samples is shown in the inverted lower panel. For the other lakes, the upper and lower panels show surface and deeper samples, respectively. Note that y-axis scales differ among panels. Source data are provided in Table S7.

Despite their shared seasonality, *Bradylimnobacteraceae* and *Kapaibacteriaceae* differed in their depth distributions (Fig. S4). *Kapaibacteriaceae* were concentrated in warm photic waters and were less frequently detected below 10 m, whereas *Bradylimnobacteraceae* extended into cold hypolimnetic waters, with detections reaching 1,200 m in Lake Tanganyika (Table S7). Consistent with this pattern, three of the four *B. soyangensis* isolates were obtained from the hypolimnion of Lake Soyang at 70 m (Table S2). Read recruitment against the isolate genomes revealed a further axis of differentiation associated with trophic status (Fig. S5). Genomes of *Bradylimnobacteraceae* and *Kapaibacteriaceae* isolates from eutrophic lakes recruited reads mainly from the summer epilimnion of other productive systems, including Lakes Mendota and Michigan and Římov Reservoir. Conversely, genomes of *Bradylimnobacteraceae* isolates from oligotrophic Lake Soyang recruited reads predominantly from nutrient-poor lakes, including Lakes Most, Zurich, and Biwa, across a broader depth range. Together, these patterns reveal niche partitioning associated with water depths and trophic status, with populations closely related to the cultured isolates recurring in geographically distant lakes.

Groundwater populations showed a contrasting spatial pattern. SingleM profiling of 808 groundwater metagenomes, including one from our sampling site, detected *Kapaibacteriales* in 145 samples, typically at relative abundances below 0.5% (Table S8 and Fig. S6). Among the detected families, UBA2268, *Kapaibacteriaceae*, and *Bradylimnobacteraceae* were the most frequent, whereas *Phreatorhabdaceae* were detected in only two metagenomes, one of which was from our sampling site. Read recruitment against individual genomes made this restricted distribution more apparent, as the genomes of the nine *Phreatorhabdaceae* isolates recruited reads only from the metagenome of their site of origin, and groundwater MAGs from other families were likewise detected only in their respective source aquifers (Fig. S7). The two groundwater *Kapaibacteriaceae* isolates did not reach the detection threshold in any metagenome, including that from their site of origin, likely because their populations were below the detection limit of read recruitment. Thus, despite the global occurrence of this order, its groundwater populations exhibited highly localized distributions within the surveyed metagenomes, consistent with a signal of subsurface endemism, although these metagenomes were geographically uneven, with 75% originating from China.

### Shared biosynthetic deficiencies and habitat-associated expansion of regulatory and mobile genes

Despite substantial differences in genome size, lake and groundwater isolates shared a core of central metabolic functions alongside extensive predicted biosynthetic deficiencies (Fig. 6a and Table S9). All 29 isolates encoded complete pathways for glycolysis, gluconeogenesis, the TCA cycle, and the non-oxidative pentose phosphate pathway, indicating a shared framework for central carbon metabolism. All isolates also encoded cytochrome *c* oxidase and F-type ATP synthase, consistent with the potential for aerobic respiration, whereas most fermentation-associated genes identified in the anaerobic thermophile *E. proteiniborus* (UBA2268) were absent. Across the global MAG collection, these fermentation-associated genes were largely restricted to the families UBA2268 and UBA4661 (Fig. S8 and Table S10).

**Fig. 6.**
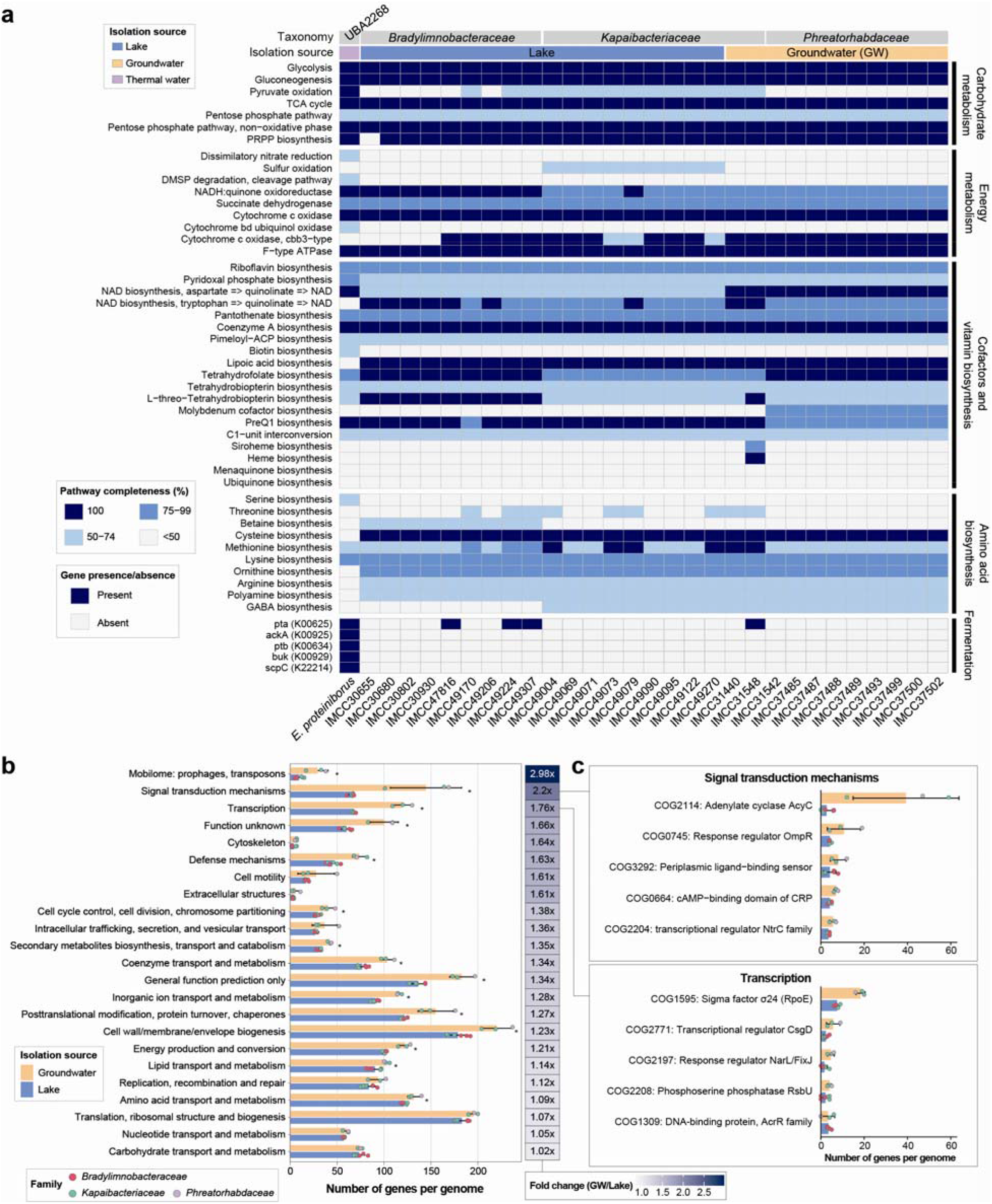
Genomic potential of *Kapaibacteriales* isolates. (a) Heatmap showing KEGG module completeness and the presence or absence of additional functional genes across isolate genomes. Taxonomy and isolation sources are indicated at the top. (b, c) Species-level comparison of Clusters of Orthologous Groups (COG) gene content between groundwater (n = 3) and lake (n = 6) species, with each species represented by a single type-strain genome. (b) Number of genes assigned to each COG functional category. The heatmap shows the fold change in mean gene number between groundwater and lake species (groundwater/lake). Differences were assessed using two-sided Wilcoxon rank-sum tests with Benjamini–Hochberg correction; asterisks indicate adjusted P < 0.05. (c) Number of genes in the five most abundant COGs within categories T (signal transduction mechanisms) and K (transcription). In (b) and (c), bars show the mean ± SD, and points represent individual species, colored by family.

Aerobic respiration requires quinones and heme, but genomic reconstruction indicated incomplete quinone^30^ biosynthesis in all isolates and incomplete heme biosynthesis pathways in all isolates except a groundwater isolate IMCC31548 (Fig. 6a and Table S9), suggesting potential dependence on external sources of these cofactors or their precursors. All isolates grew in bovine catalase-supplemented medium without separate addition of either cofactor (Table S1). However, supplementation with hemin, menaquinone-4, or both did not support the growth of a representative strain IMCC31440 in the absence of the catalase preparation under the conditions tested (Fig. S9), although this isolate encodes a gene for KatG, a heme-dependent catalase (Table S9). Peroxide removal, cofactor supply, including quinones or quinone precursors, and other undefined components of the catalase preparation therefore remain unresolved explanations for its growth-promoting effect.

Biosynthetic deficiencies also extended to amino acids and vitamins. Only a limited set of amino acid biosynthesis pathways reached ≥50% completeness, including those for cysteine, methionine, lysine, ornithine, and arginine, whereas the shikimate pathway and the pathways for branched-chain amino acids and histidine were absent or largely incomplete in all isolates (Table S9). Among the vitamin and cofactor biosynthesis pathways examined, those for riboflavin, NAD, pantothenate, pyridoxal phosphate, and tetrahydrofolate reached the same threshold, whereas those for thiamine, biotin, and cobalamin did not. These predicted deficiencies were shared between compact lake genomes and larger groundwater genomes and were also widespread among MAGs from diverse habitats, with exceptions in a small number of UBA4661 genomes (Fig. S8).

Against this background of shared biosynthetic deficiencies, groundwater isolates carried more genes than lake isolates in the COG categories of mobilome (2.98-fold), signal transduction (2.2-fold), and transcription (1.76-fold) (Fig. 6b and Table S11). The larger mobilome was accompanied by prophages, which accounted for 1.1–4.5% of groundwater genomes but were not detected in lake genomes (Fig. S10 and Table S12). These prophage regions contained putative antibiotic resistance genes, genes encoding phosphoadenosine phosphosulfate (PAPS) reductase involved in sulfur assimilation, and components of the DISARM defense system (Table S13), suggesting that integrated prophages may contribute functional capacity beyond the mobile elements themselves. The larger signal transduction and transcription repertoires were associated with elevated paralog content. The proportion of genes classified as paralogs was more than twice as high in groundwater genomes as in lake genomes (10.8% vs. 4.9%; Fig. S11), and paralogs accounted for much of the difference in gene counts within both categories. Adenylate cyclase genes averaged 39.3 copies per groundwater genome compared with 2.7 per lake genome (Fig. 6c), and genes encoding the sigma factor σ^24^ (RpoE) showed a similar but smaller difference (18.3 vs. 7.5 copies).

Several functional systems showed additional habitat-or family-associated distributions. All groundwater isolates encoded the nitrous oxide reductase gene cluster *nosZDFYL*, which was absent from lake isolates (Fig. S12 and Table S9). *Phreatorhabdaceae* isolates additionally encoded the nitrate reductase genes *narGH*, but none of the *nosZ*-containing isolates encoded nitrite reductase genes (*nirK* or *nirS*), suggesting a potential for N_2_O reduction without complete denitrification^31^. In contrast, all lake isolates encoded microbial rhodopsins together with the genes required for retinal biosynthesis (*crtE, crtB*, *crtI*, *crtY*, and *blh*), the last two genes of which were absent from groundwater isolates. Other differences aligned with family membership. Zinc/manganese/iron transporters (*troABCD*) were found only in *Bradylimnobacteraceae*, whereas complete or near-complete flagellar gene sets were restricted to *Phreatorhabdaceae*, consistent with the flagella observed exclusively in these isolates (Fig. S3). *Phreatorhabdaceae* isolates also encoded CRISPR–Cas systems and a type VI secretion system, suggesting potential for antiviral defense and contact-dependent interactions, respectively (Fig. S12 and Tables S14– S15).

## Discussion

The 29 axenic isolates obtained here are the first cultivated representatives of *Kapaibacteriales* from non-thermal environments and establish a taxonomic and experimental foundation for this order, which was previously known almost exclusively from environmental sequences (Fig. 1). By combining these cultures with global metagenomic surveys, we identified two contrasting lifestyles within the order. Lake populations were widespread, recurrently enriched in the summer epilimnion, and carried relatively compact genomes, whereas groundwater populations were geographically restricted and carried larger genomes enriched in regulatory, signaling, and mobile genes (Figs. 2, 4–6). Despite these differences, both populations broadly shared predicted deficiencies in amino acid, vitamin, and respiratory cofactor biosynthesis. The central implication of these findings is that greater coding capacity can support a broader regulatory and accessory repertoire without a corresponding increase in biosynthetic self-sufficiency.

Genome reduction is common among oligotrophic aquatic bacteria^32,33^, and the larger genomes of groundwater populations in oligotrophic environments therefore indicate that resource scarcity alone is not sufficient to predict genome architecture in this order. A similar groundwater–lake contrast in genome size has been reported for *Methylotenera*^34^, so the pattern is not unique to *Kapaibacteriales*. It is nevertheless not universal, as *Patescibacteriota* retain reduced genomes in both habitats^14,35^. Studies of other aquifer systems offer further context for large genomes in nutrient-poor groundwater. A large-scale survey in China reported increasing average genome size with depth^15^, and a study in Northern Europe identified a localized peak in estimated genome size in the deepest, most hydrologically isolated borehole, although smaller genomes predominated elsewhere^16^. Together, these observations suggest that genome expansion in groundwater bacteria depends on lineage and local hydrogeological conditions rather than on resource availability alone.

The functional composition of the genomes provided insight into the nature of this contrast. Groundwater genomes carried more genes for signal transduction and transcriptional regulation, with paralog expansion particularly pronounced in genes encoding adenylate cyclases and RpoE-type sigma factors (Fig. 6b, c). Some increase in regulatory genes is expected from genome-size scaling^4,5^, but the concentration of this difference in a few paralogous gene families points to an expanded capacity to sense and respond to environmental conditions rather than a general expansion of metabolic function. One interpretation is that such regulatory capacity supports exploitation of the spatially heterogeneous resources and episodic nutrient inputs that often characterize groundwater systems^13^. This remains a hypothesis, however, because the *in situ* resource dynamics experienced by these populations were not measured in this study. Prophages, which were absent from lake genomes, also contributed to the larger groundwater genomes (Fig. S10), suggesting that both gene duplication and horizontal acquisition contributed to the genome-size difference. The highly localized distribution of groundwater populations, in contrast to the cosmopolitan distribution of lake populations, further suggests that limited dispersal in the subsurface may allow lineage-specific features, including prophages and CRISPR–Cas systems, to accumulate locally.

Lake populations of *Kapaibacteriales*, by contrast, appear to be closely coupled to phytoplankton productivity. Their recurrent summer enrichment, co-occurrence with cyanobacteria, and highest relative abundances in phytoplankton-associated samples, including cyanobacterial cultures, support this point (Figs. 4 and 5). Conversely, neither *Bradylimnobacteraceae* nor *Kapaibacteriaceae* were detected in dystrophic bog lakes, where dissolved organic matter is largely of terrestrial origin, suggesting that their occurrence depends on autochthonous production (Table S7). All lake isolates also encoded microbial rhodopsins together with the genes required for retinal biosynthesis, some of which were not found in groundwater isolates, suggesting that light may supplement heterotrophic energy acquisition in the photic zone. Within this shared lifestyle, depth and trophic status partitioned populations, with *Bradylimnobacteraceae* extending into cold hypolimnia and related populations recurring in geographically distant lakes of similar trophic status (Figs. S4 and S5). This combination of compact genomes, photoheterotrophic potential, and extensive predicted auxotrophy resembles that of other abundant freshwater lineages such as *Nanopelagicales^11,12,32^*, suggesting a convergent strategy among freshwater bacteria that exploit phytoplankton-derived resources.

The persistence of predicted biosynthetic deficiencies despite this genomic divergence was particularly intriguing (Fig. 6a). A previous large-scale survey associated smaller prokaryotic genomes with more extensive amino acid auxotrophy^7^, yet the larger groundwater *Kapaibacteriales* genomes did not contain more complete versions of the pathways missing from lake genomes, including the biosynthetic pathways for branched-chain amino acids and vitamins such as thiamine, biotin, and cobalamin. Genome size and predicted biosynthetic self-sufficiency therefore appear to vary independently within this order, consistent with a recent analysis identifying phylogeny as a stronger predictor of amino acid auxotrophy than environmental context^36^. Adaptive loss of functions supplied by surrounding organisms, as proposed by the BQH, offers one explanation for the origin of such dependencies, although the present data do not establish whether the losses were ancestral or recurrent, whether they were selectively favored, or whether the required compounds are obtained through active exchange or from environmental pools.

The cofactor deficiencies are of particular interest in light of the catalase requirement of the isolates. All isolates lacked a complete quinone biosynthesis pathway, and all except IMCC31548 also lacked a complete heme biosynthesis pathway, suggesting a requirement for exogenous cofactors or their precursors. Such cofactor dependencies have been demonstrated for quinones in gut bacteria^37^ and for heme in abundant aquatic lineages^22^. In *Planktophila rubra* IMCC25003 and *Planktophila aquatilis* IMCC26103 of *Nanopelagicales^22^*, for example, a comparable requirement for catalase was traced to heme auxotrophy, with the bovine catalase preparation serving as a heme source, and strain IMCC26103 additionally required riboflavin present as an impurity in the preparation. By analogy, the bovine catalase used here may supply not only heme but also quinones or quinone precursors. Although hemin and MK-4 together did not support the growth of IMCC31440 without catalase (Fig. S9), this result does not exclude a quinone requirement, because the isolates may require quinones with different side chains. Identifying the components of the catalase preparation that support growth, for example, through supplementation with alternative quinones and their precursors, would clarify whether combined heme and quinone auxotrophy underlies the cultivation requirements of this order.

Cultivation added an organismal perspective that refines the interpretation of these genomic and ecological comparisons. Temperature responses were partly consistent with the thermal regimes of the contrasting source environments, with groundwater isolates growing better at 4 °C and a lake isolate tolerating 37 °C, but all isolates grew fastest at 25–30 °C, well above *in situ* groundwater temperatures (Fig. 3). Moreover, two *Phreatorhabdaceae* isolates sharing >99.9% ANI differed substantially in growth rate and yield, indicating that high genomic similarity does not imply physiological equivalence. Several limitations should also be considered. Groundwater isolates were obtained from a single aquifer, and the families recovered from lakes and groundwater only partly overlapped, so habitat and phylogeny are not fully separable in comparisons based solely on the isolates. In addition, the groundwater metagenomes surveyed were geographically uneven, and biosynthetic deficiencies were inferred from gene content and remain to be confirmed experimentally.

Together, these findings document habitat-associated genomic differentiation alongside shared biosynthetic constraints within a single cultivated lineage. They suggest that ecological diversification can involve substantial changes in environmental responsiveness, energy acquisition, and accessory functions while leaving many biosynthetic dependencies in place, even where coding capacity increases. The cultures and taxonomic framework established here provide a tractable system for linking these genomic patterns to the resources, interactions, and physiological responses that sustain *Kapaibacteriales* across contrasting environments.

## Methods

### Sample collection, isolation, and culture

A total of 17 high-throughput cultivation campaigns based on dilution-to-extinction culturing were conducted between 2017 and 2023 at four freshwater sampling sites in the Republic of Korea, of which 10 yielded *Kapaibacteriales* isolates (Fig. 1a and Tables S2-S3). Water samples were collected from the surface and selected depths of Lake Soyang (1 and 70 m), Lake Paldang (1, 10, and 20 m), and Lake Daechung (1 m), and groundwater was collected from a 120-m-deep well in Yanggu after more than 200 L of water had been purged. Physicochemical parameters, including temperature, pH, conductivity, and dissolved oxygen, were measured *in situ* using a YSI 556 MPS multiprobe (YSI Inc., Yellow Springs, OH, USA). For chlorophyll *a* measurement, water samples were filtered through GF/F filters (Pall, Port Washington, NY, USA), and the retained biomass was extracted in 90% acetone and quantified using a UV-2600 UV-VIS spectrophotometer (Shimadzu, Kyoto, Japan). Dissolved nutrient concentrations, including nitrate, nitrite, ammonium, phosphate, and silicate, were measured from water filtered through a 0.45-µm mixed cellulose ester membrane (Advantec, Tokyo, Japan), using a HACH DR/2010 spectrophotometer (Hach, Loveland, CO, USA). Total organic carbon, dissolved organic carbon, total phosphorus, and total nitrogen were quantified at the National Instrumentation Center for Environmental Management (NICEM, Seoul, Republic of Korea) following Standard Methods for the Examination of Water and Wastewater^38^.

To obtain cultures of diverse freshwater bacteria, we followed previously established protocols that mimic the physicochemical conditions of the corresponding natural environments^39^. Briefly, a low-nutrient freshwater medium (LNFM; Table S1) was prepared using natural water collected on-site. The water was filtered through 0.22-μm membrane filters, autoclaved for 1.5 h, aerated for 6 h, and amended with carbon sources, amino acids, vitamins, and bovine catalase. Total prokaryotic cell numbers in unfiltered water samples were determined using a Guava EasyCyte Plus flow cytometer (Millipore, Billerica, MA, USA) after staining with SYBR Green I (Invitrogen, Carlsbad, CA, USA). Cells were inoculated into the LNFM at a final concentration of 0.5–10 cells mL^-1^, and 1 mL aliquots were dispensed into 48-well microtiter plates (BD Falcon, Franklin Lakes, NJ, USA). Following incubation at 18–20 °C for up to one month, a subsample from each well was stained with SYBR Green I (Invitrogen) and analyzed via flow cytometry. For wells exhibiting growth (>5 × 10^4^ or 10^5^ cells mL ¹, depending on the campaign), 400-μL aliquots were cryopreserved in 10% (v/v) glycerol suspension at −80 °C. For the taxonomic identification of the cultures, cell lysates were prepared either via three freeze– thaw cycles or by using InstaGene Matrix (Bio-Rad Laboratories, Hercules, CA, USA) according to the manufacturer’s protocols. The 16S rRNA gene was amplified using the universal primers 27F and 1492R. For samples that failed initial PCR amplification, a nested PCR was subsequently performed using the primers 27F and 1406R. The resulting amplicons were subjected to Sanger sequencing. The generated sequences were then classified against the SILVA SSURef NR99 database (release 138)^40^ using the *classify.seqs* function in Mothur v1.39.5^41^. Cultures affiliated with *Kapaibacteriales* were selected for further analyses.

### Whole-genome sequencing, assembly, and taxonomy of isolates

Strain IMCC31440, one of the fastest-growing isolates, was cultured in 1 L of LNFM and harvested by centrifugation at 10,000 × *g* for 10 min at 4 °C. Genomic DNA was extracted from the cell pellet using the DNeasy Blood & Tissue Kit (Qiagen, Hilden, Germany). For the remaining isolates, genomic DNA was amplified directly from glycerol stocks via multiple displacement amplification using the REPLI-g Single Cell Kit (Qiagen, Hilden, Germany). Whole-genome sequencing libraries were prepared and sequenced by DNA Link Inc. (Seoul, Republic of Korea) on the NovaSeq X Plus platform (Illumina, San Diego, CA, USA). Raw reads were quality-filtered to remove low-quality bases (Phred score < 10), adapter sequences, and PhiX contaminants using the bbduk.sh script from the BBMap package (https://sourceforge.net/projects/bbmap/). The filtered reads were assembled using SPAdes v3.15.4^42^ with k-mer sizes of 21, 33, 55, 77, 99, and 127. Genome completeness and contamination were evaluated using CheckM2 v1.1.0^43^. Pairwise average nucleotide identity (ANI) among isolate genomes was calculated using FastANI v1.34^44^, and genomes sharing ≥95% ANI were assigned to the same genomic species. Taxonomic assignment for each genome was performed using GTDB-Tk v2.4.1 based on the GTDB r226 database^45^. Names of novel taxa were proposed in accordance with the SeqCode^46^, with isolate genomes designated as nomenclatural types.

### Global MAG dataset and phylogenomic analysis

Public genome data were retrieved from the GlobDB R226^47^. MAGs affiliated with the order *Kapaibacteriales*, exhibiting completeness ≥90% and contamination ≤5% as estimated by CheckM2 v1.1.0^43^, were selected for analysis (*n* = 248; Table S5). Maximum-likelihood phylogenomic trees were constructed using the isolate genomes and the MAGs. A core set of 120 conserved bacterial marker proteins was identified, aligned, and concatenated using the identify and align workflows of GTDB-Tk v2.4.1^45^. The resulting sequence alignment was subsequently used as input for IQ-TREE2 v2.2.0^48^ with the following parameters: --perturb 0.2 --nstop 500 -B 1000 -m TEST --alrt 1000. The Q.yeast+I+G4 model was selected as the best-fitting evolutionary model by ModelFinder based on the Bayesian Information Criterion. The genome of *Flavobacterium johnsoniae* was used as the outgroup to root the tree. The phylogenomic tree was visualized and annotated using the interactive Tree Of Life (iTOL) v6^49^.

### Genome annotation and metabolic pathway analysis

All isolate genomes and selected high-quality MAGs were used for functional and metabolic pathway analyses. Open reading frame (ORF) prediction and functional annotation were performed in Anvi’o v8^50^. ORFs were predicted and translated using Prodigal^51^ during the creation of the contig database via the *anvi-gen-contigs-database* function. Functional annotations were assigned by querying the translated protein-coding sequences against multiple curated databases, including the COG (*anvi-run-ncbi-cogs*) and KOfam (*anvi-run-kegg-kofams*) databases. Additionally, KEGG Orthology (KO) assignments were supplemented using BlastKOALA^52^, and the KO terms obtained from KOfam and BlastKOALA were manually integrated. Metabolic module completeness was evaluated using the kegg-pathways-completeness tool under the MGnify platform^53^. Flagellar and type VI secretion systems (T6SS) were predicted using TXSScan in MacSyFinder v2.1.6^54^, and CRISPR-Cas systems were identified using CCTyper v1.5.0^55^. Orthogroups were inferred using using OrthoFinder v3.1.3^56^. For each genome, paralog counts were calculated by summing the number of additional gene copies beyond the first copy within each orthogroup. Prophage sequences were identified using geNomad v1.11.1^57^, and ORFs within prophage regions were predicted using Pharokka v1.7.4^58^, followed by functional annotation using Phold v0.2.0^59^.

### Groundwater metagenomic sequencing

For metagenomic sequencing, 8 L of groundwater collected from the GW46 site on November 2, 2018, was initially filtered through a 3 µm cellulose acetate membrane filter (Advantec) to remove coarse particles and larger organisms, and the filtrate was passed through a 0.22-µm polyethersulfone membrane filter (Pall) to collect microbial biomass. The membrane filters were stored at −80 °C until further processing. Genomic DNA was extracted using the DNeasy PowerWater DNA Isolation Kit (Qiagen), and the DNA concentration was quantified using a Qubit fluorometer (Invitrogen). Sequencing library preparation and high-throughput sequencing were performed by CJ Bioscience (Seoul, Republic of Korea) on the NovaSeq 6000 platform (Illumina, San Diego, CA, USA) to generate 10–15 Gb of data with 2 × 150 bp paired-end reads.

### Global distribution analysis

The global distribution of *Kapaibacteriales* was assessed using the Sandpiper database v1.1.9^29^. Profiles assigned to *Kapaibacteriales* were retrieved along with sample metadata, including habitat descriptions and geographic coordinates. To examine finer-scale distributions in freshwater and groundwater environments, a total of 2,067 publicly available freshwater metagenomes and 808 groundwater metagenomes were analyzed. Taxonomic profiling of these metagenomes was performed using SingleM v0.20.3^29^. Subsequently, metagenome fragment recruitment was conducted on this same metagenomic dataset. Before fragment recruitment, 16S rRNA genes in the isolate genomes were masked to prevent false-positive mappings resulting from highly conserved sequences. Metagenomic reads were mapped against the masked genomes using CoverM^60^ with a minimum read identity threshold of 97% *(--min-read-percent-identity 97*). A genome was considered present in a metagenomic sample when its covered fraction exceeded 0.5 (i.e., more than 50% of the genome length was covered). Relative abundance was calculated as the number of mapped reads per kilobase of genome per million metagenomic reads (RPKM). Finally, abundance profiles were visualized as heatmaps using the R package pheatmap v1.0.12 (https://github.com/raivokolde/pheatmap).

### Growth experiments

Five representative isolates were selected to evaluate their growth characteristics. A 1% (v/v) inoculum from an exponential-phase culture was transferred into 20 mL of fresh LNFM, resulting in initial cell densities of approximately 10^3^ (IMCC49307) to 10^5^ cells mL^-1^. To ensure consistency across strains, LNFM prepared with filtered water from Lake Soyang was used as the basal medium for all strains. Cellular growth was tested at seven temperatures (4, 10, 15, 20, 25, 30, and 37 °C) in triplicate. Cell density was monitored every two to three days for up to 24 days. Culture purity was verified by 16S rRNA gene sequencing both before inoculation and at the end of each experiment.

Cell concentrations were quantified via flow cytometry using a Guava EasyCyte Plus system (Millipore), except for strain IMCC49307. Because the cell densities of IMCC49307 fell below the lower linear detection range of the flow cytometer, its cell abundance was determined by direct microscopic enumeration. Cells of this strain were stained with SYBR Green I (Invitrogen) and counted under an epifluorescence microscope (Nikon Eclipse 80i, Nikon, Tokyo, Japan). Specific growth rates (day^-1^) were estimated from time-course cell density data using the easylinear method implemented in the R package growthrates v. 0.8.5^61^.

To determine whether hemin or menaquinone-4 could substitute for catalase in supporting growth, IMCC31440 was cultured in catalase-free LNFM under five conditions: no supplementation, bovine liver catalase (C1345; Sigma-Aldrich, St. Louis, MO, USA), hemin (Sigma-Aldrich), menaquinone-4 (MK-4; Sigma-Aldrich), or hemin and MK-4 in combination. Supplements were added at the final concentrations indicated in Fig. S9. All treatments were performed in triplicate and incubated at 25 °C for 5 days. Cell densities were measured daily by flow cytometry to compare growth responses among treatments.

### Electron microscopy

The five selected isolates were cultivated to the stationary phase at 25 °C. Scanning electron microscopy (SEM) was performed according to a previously described protocol^62^ with minor modifications. Cultures were fixed with 0.2% (v/v) glutaraldehyde (Merck, Darmstadt, Germany) in 0.1 M sodium cacodylate buffer (Acros Organics, Geel, Belgium) at 4 °C for 1 h. The fixed cells were collected on 0.22-µm polycarbonate membrane filters (MilliporeSigma, Burlington, MA, USA), dehydrated through a graded ethanol series (30% to 100%, 10 min per step), and chemically dried using sequential mixtures of absolute ethanol and hexamethyldisilazane (HMDS; Sigma-Aldrich, St. Louis, MO, USA) (2:1, 1:1, and 1:2), followed by 100% HMDS treatment. The dried filters were mounted on aluminum stubs, carbon-coated, and imaged using a Hitachi S-4300 SEM (Hitachi, Osaka, Japan).

For transmission electron microscopy (TEM), cells were harvested from 1 L cultures by centrifugation at 10,000 × *g* for 90 min at 4 °C and resuspended in 2 mL of sterile LNFM. The suspension was applied to Formvar/carbon-coated copper grids (Ted Pella, Redding, CA, USA) and allowed to adsorb for 3 min. After excess liquid was blotted with filter paper (No. 2, Advantec), the grids were negatively stained with 0.5% (w/v) uranyl acetate (Merck) for 3 min in the dark. The grids were air-dried and imaged using a Talos L120C TEM (Thermo Fisher Scientific, Waltham, MA, USA) at an accelerating voltage of 120 kV.

### Statistical analysis

All statistical analyses were performed in R v4.4.2. Associations were assessed using Spearman’s rank correlation, and differences between groups were evaluated using two-sided Wilcoxon rank-sum tests. Wilcoxon test *P*-values were adjusted for multiple comparisons using the Benjamini–Hochberg false discovery rate (FDR) procedure.

## Data availability

The genome sequences of the 29 isolates and the groundwater metagenome generated in this study have been deposited in the European Nucleotide Archive (ENA) under BioProject accessions PRJEB121484 and PRJEB128671, respectively. BioSample accessions of the isolate genomes are listed in Table S2. Names of the novel taxa proposed in this study have been submitted to the SeqCode Registry and will become validly published upon validation of the list following publication of this article.

