## Supplementary_information for "Cultivated *Kapaibacteriales* reveal contrasting lake and groundwater lifestyles built on shared metabolic dependencies"

**This file includes:**

Supplementary Notes
Supplementary Figs. S1– S12

### Supplementary Notes

#### Descriptions of new taxa

This note provides descriptions and etymologies for the two novel families, three novel genera, and nine novel species proposed in this study, including one novel species of the genus *Kapaibacterium*. In addition, emended description of the family *Kapaibacteriaceae* and the genus *Kapaibacterium* are provided to include phenotypic properties of the first cultivated representative. Names were proposed in accordance with the SeqCode and have been submitted to the SeqCode Registry.

#### Emended description of the family *Kapaibacteriaceae*

The description of the family is as given by Chuvochina *et al*. (2023), with the following additions. The family was previously known only from metagenome-assembled genomes, which originate mainly from freshwater and wastewater environments. Cultivated representatives belong to the genera *Kapaibacterium* and *Dulcisphaera*. These genera are available exclusively as dilution-to-extinction cultures, obtained from a freshwater lake and groundwater. Cells are coccoid and grow in liquid low-nutrient freshwater medium supplemented with bovine catalase.

Registry URL: https://seqco.de/i:31935

#### Emended description of the genus *Kapaibacterium*

The description of the genus is as given by Kantor *et al*. (2015), with the following additions. The genus was previously known only from metagenome-assembled genomes. Cells are coccoid. Does not form colonies on agar surface. The first cultivated representative was obtained from groundwater by dilution-to-extinction cultivation and grows aerobically in liquid low-nutrient freshwater medium supplemented with bovine catalase. Genome sizes of the cultivated representative and of the metagenome-assembled genomes assigned to the genus are 2.5–3.4 Mbp, with DNA G+C contents of 53–59%. *Kapaibacterium aquifericola* is the first and currently the only cultivated species of the genus. The nomenclatural type remains the genome sequence of *Kapaibacterium thiocyanatum*, a metagenome-assembled genome.

Registry URL: https://seqco.de/i:31934

#### Description of *Kapaibacterium aquifericola* sp. nov.

*Kapaibacterium aquifericola* (a.qui.fe.ri.co'la. L. masc. adj. *aquifer*, water-bearing; L. suff. *-cola* (from L. n. *incola*), inhabitant, dweller; N.L. n. *aquifericola*, an inhabitant of an aquifer).

The type strain is IMCC31440, obtained from groundwater of the Yanggu aquifer, Republic of Korea, by dilution-to-extinction cultivation in liquid low-nutrient freshwater medium supplemented with bovine catalase. Cells of the type strain are coccoid, approximately 0.3–0.5 μm in diameter, as observed by electron microscopy. Does not form colonies on agar surface. Growth occurs at 4–37 °C (optimum 25–30 °C), with the highest specific growth rate (approximately 1.5 day$^{-1}$) and maximum cell density (approximately $1.0 \times 10^7$ cells mL$^{-1}$) at 25–30 °C. Growth requires the bovine catalase preparation, and supplementation with hemin, menaquinone-4, or both does not restore growth in its absence. The genome encodes complete pathways for central carbon metabolism and components of aerobic respiration, as well as nitrous oxide reductase, whereas predicted biosynthetic pathways for quinones and heme are incomplete. The species is the first cultivated representative of the genus. The draft genome assembly of IMCC31440 comprises 3,377,527 bp, with a DNA G+C content of 56.75% and 2,804 predicted genes. The species belongs to g__*Kapaibacterium* but is not assigned to any species in GTDB (R226). The nomenclatural type is the genome sequence of strain IMCC31440 (ENA WGS accession number CFWNPL01).

Registry URL: https://seqco.de/i:57155

**Description of *Dulcisphaera* gen. nov.**

*Dulcisphaera* (Dul.ci.sphae'ra. L. masc. adj. *dulcis*, sweet; L. fem. n. *sphaera*, a ball, sphere; N.L. fem. n. *Dulcisphaera*, a sphere from fresh water).

Cells are coccoid. Does not form colonies on agar surface. Cultivated representatives have been obtained from a freshwater lake and groundwater in the Republic of Korea by dilution-to-extinction cultivation and grow in liquid low-nutrient freshwater medium supplemented with bovine catalase. Genome sizes are 2.4–3.4 Mbp, with DNA G+C contents of 51.3–54.6%. The genus corresponds to g__UBA10438 within the family *Kapaibacteriaceae* in GTDB (R226). The type species is *Dulcisphaera aestivalis*.

Registry URL: https://seqco.de/i:57156

**Description of *Dulcisphaera aestivalis* sp. nov.**

*Dulcisphaera aestivalis* (ae.sti.va'lis. L. fem. adj. *aestivalis*, of or belonging to summer).

The type strain is IMCC49270, obtained from Lake Daechung, Republic of Korea, by dilution-to-extinction cultivation in liquid low-nutrient freshwater medium supplemented with bovine catalase. Cells of the type strain are coccoid, approximately 0.3–0.5 μm in diameter, as observed by electron microscopy. Growth occurs at 10–37 °C (optimum 25 °C), with little growth at 4 °C. The highest specific growth rate (approximately 1.5 day$^{-1}$) and maximum cell density (approximately $1.4 \times 10^7$ cells mL$^{-1}$) occur at 25 °C, and similar values are observed at 37 °C. Additional cultured isolates assigned to the same species are IMCC49069, IMCC49071, IMCC49090, IMCC49095, and IMCC49122, obtained from the same lake. The genome encodes complete pathways for central carbon metabolism and components of aerobic respiration, as well as a proton-pumping rhodopsin, whereas predicted biosynthetic pathways for quinones and heme are incomplete. The genome shares 81–84% ANI with those of the type strains of *D. daechungensis* and *D. receptaculi*. The draft genome assembly of IMCC49270 comprises 2,456,917 bp, with a DNA G+C content of 54.56% and 2,070 predicted genes. The species belongs to g__UBA10438 but is not assigned to any species in GTDB (R226). The nomenclatural type is the genome sequence of strain IMCC49270 (ENA WGS accession number CFWNPE01).

Registry URL: https://seqco.de/i:57157

**Description of *Dulcisphaera phreaticola* sp. nov.**

*Dulcisphaera phreaticola* (phre.a.ti'co.la. Gr. neut. n. *phrear,* a well; L. suff. *-cola* (from L. n. *incola*), inhabitant, dweller; N.L. fem. n. *phreaticola*, a dweller of groundwater).

The type strain is IMCC31548, obtained from groundwater of the Yanggu aquifer, Republic of Korea, by dilution-to-extinction cultivation in liquid low-nutrient freshwater medium supplemented with bovine catalase. Growth of the type strain was observed at 20 °C, the temperature used for isolation; other cultural properties have not been determined. The genome encodes complete pathways for central carbon metabolism, heme biosynthesis, and components of aerobic respiration, as well as nitrous oxide reductase, whereas the predicted quinone biosynthesis pathway is incomplete. Unlike the other species of the genus, the genome encodes a complete heme biosynthesis pathway and lacks rhodopsin, and both the genome size (approximately 3.4 Mbp) and DNA G+C content (approximately 51%) differ from those of the other species (approximately 2.5 Mbp and 54–55%, respectively). The draft genome assembly of IMCC31548 comprises 3,396,867 bp, with a DNA G+C content of 51.27% and 2,830 predicted genes. The species corresponds to s__UBA10438 sp937870915 in GTDB (R226). The nomenclatural type is the genome sequence of strain IMCC31548 (ENA WGS accession number CFWNPH01).

Registry URL: https://seqco.de/i:57158

**Description of *Dulcisphaera daechungensis* sp. nov.**

*Dulcisphaera daechungensis* (dae.chung.en'sis. N.L. fem. adj. *daechungensis*, pertaining to Lake Daechung, an artificial reservoir from which the type strain was isolated).

The type strain is IMCC49004, obtained from Lake Daechung, Republic of Korea, by dilution-to-extinction cultivation in liquid low-nutrient freshwater medium supplemented with bovine catalase. Growth of the type strain was observed at 18 °C, the temperature used for isolation; other cultural properties have not been determined. Another cultured isolate assigned to the same species is IMCC49073, obtained from the same lake. The genome encodes complete pathways for central carbon metabolism and components of aerobic respiration, as well as a proton-pumping rhodopsin, whereas predicted biosynthetic pathways for quinones and heme are incomplete. The genome shares approximately 81–82% ANI with those of the type strains of *D. aestivalis* and *D. receptaculi*. The draft genome assembly of IMCC49004 comprises 2,451,049 bp, with a DNA G+C content of 53.95% and 2,043 predicted genes. The species corresponds to s__UBA10438 sp030830285 in GTDB (R226). The nomenclatural type is the genome sequence of strain IMCC49004 (ENA WGS accession number CFWNPF01).

Registry URL: https://seqco.de/i:57159

**Description of *Dulcisphaera receptaculi* sp. nov.**

*Dulcisphaera receptaculi* (re.cep.ta'cu.li. L. gen. n. *receptaculi*, of a reservoir).

The type strain is IMCC49079, obtained from Lake Daechung, Republic of Korea, by dilution-to-extinction cultivation in liquid low-nutrient freshwater medium supplemented with bovine catalase. Growth of the type strain was observed at 18 °C, the temperature used for isolation; other cultural properties have not been determined. The genome encodes complete pathways for central carbon metabolism and components of aerobic respiration, as well as a proton-pumping rhodopsin, whereas predicted biosynthetic pathways for quinones and heme are incomplete. The genome shares 81–84% ANI with those of the type strains of *D. aestivalis* and *D. daechungensis*. The draft genome assembly of IMCC49079 comprises 2,504,300 bp, with a DNA G+C content of 54.30% and 2,083 predicted genes. The species corresponds to s__UBA10438 sp903939735 in GTDB (R226). The nomenclatural type is the genome sequence of strain IMCC49079 (ENA WGS accession number CFWNPR01).

Registry URL: https://seqco.de/i:57160

**Description of *Bradylimnobacteraceae* fam. nov.**

*Bradylimnobacteraceae* (Bra.dy.lim.no.bac.te.ra'ce.ae. N.L. masc. n. *Bradylimnobacter*, type genus of the family; L. fem. pl. n. suff. *-aceae*, ending to denote a family; N.L. fem. pl. n. *Bradylimnobacteraceae*, the *Bradylimnobacter* family).

The family was previously known only from metagenome-assembled genomes and is represented in GTDB (R226) as the family f__UBA961, named after genome UBA961, which was reconstructed from public metagenomes. These genomes originate mainly from freshwater and wastewater environments. Cultivated representatives first obtained are available exclusively as dilution-to-extinction cultures. In GTDB (R226), the family comprises four genera, of which *Bradylimnobacter* corresponds to g__UBA961. The description of the family is otherwise the same as that of the genus *Bradylimnobacter*. The type genus is *Bradylimnobacter*.

Registry URL: https://seqco.de/i:57131

**Description of *Bradylimnobacter* gen. nov.**

*Bradylimnobacter* (Bra.dy.lim.no.bac'ter. Gr. masc. adj. *bradys*, slow; Gr. fem. n. *limnē*, a lake; N.L. masc. n. *bacter*, a rod; N.L. masc. n. *Bradylimnobacter*, a slow-growing rod from a lake).

Cells are small rods and grow slowly. Does not form colonies on agar surface. Cultivated representatives have been obtained from freshwater lakes in the Republic of Korea by dilution-to-extinction cultivation and grow in liquid low-nutrient freshwater medium supplemented with bovine catalase. Genome sizes are 2.4–2.6 Mbp, with DNA G+C contents of 33.0–40.1%. The genus corresponds to g__UBA961 in GTDB (R226). The type species is *Bradylimnobacter exiguus*.

Registry URL: https://seqco.de/i:57148

**Description of *Bradylimnobacter exiguus* sp. nov.**

*Bradylimnobacter exiguus* (e.xi'gu.us. L. masc. adj. *exiguus*, scanty, meagre, small in amount, referring to the very low biomass yield of the type strain).

The type strain is IMCC49307, obtained from Lake Daechung, Republic of Korea, by dilution-to-extinction cultivation in liquid low-nutrient freshwater medium supplemented with bovine catalase. Cells of the type strain are small rods, approximately 0.2 × 0.4 µm, as observed by electron microscopy. Growth occurs at 10–30 °C (optimum 25 °C) and is slow, with the highest specific growth rate (approximately 0.4 $day^{-1}$) and maximum cell density (approximately $7.5 \times 10^4$ cells $mL^{-1}$) at 25 °C. Additional cultured isolates assigned to the same species are IMCC49170 and IMCC49224, obtained from the same lake. The genome encodes complete pathways for central carbon metabolism and components of aerobic respiration, as well as a proton-pumping rhodopsin, whereas predicted biosynthetic pathways for quinones and heme are incomplete. The DNA G+C content (approximately 40%) is higher than those of the other species of the genus (approximately 33 mol%). The draft genome assembly of IMCC49307 comprises 2,535,771 bp, with a DNA G+C content of 40.01%, 2,090 predicted genes. The species corresponds to s__UBA961 sp002293005 in GTDB (R226). The nomenclatural type is the genome sequence of strain IMCC49307 (ENA WGS accession number CFWNOT01).

Registry URL: https://seqco.de/i:57149

**Description of *Bradylimnobacter soyangensis* sp. nov.**

*Bradylimnobacter soyangensis* (so.yang.en'sis. N.L. masc. adj. *soyangensis*, pertaining to Lake Soyang, an artificial reservoir from which the type strain was isolated).

The type strain is IMCC30655, obtained from Lake Soyang, Republic of Korea, by dilution-to-extinction cultivation in liquid low-nutrient freshwater medium supplemented with bovine catalase.

Growth of the type strain was observed at 20 °C, the temperature used for isolation; other cultural properties have not been determined. Additional cultured isolates assigned to the same species are IMCC30680, IMCC30802, and IMCC30930, obtained from the same lake. The genome encodes complete pathways for central carbon metabolism and components of aerobic respiration, as well as a proton-pumping rhodopsin, whereas predicted biosynthetic pathways for quinones and heme are incomplete. The genome shares approximately 91% ANI with that of the type strain of *B. lacuum*, and the DNA G+C content (approximately 33%) is lower than that of *B. exiguus* (approximately 40%). The draft genome assembly of IMCC30655 comprises 2,485,935 bp, with a DNA G+C content of 33.18% and 2,057 predicted genes. The species corresponds to s__UBA961 sp014191195 in GTDB (R226). The nomenclatural type is the genome sequence of strain IMCC30655 (ENA WGS accession number CFWNOU01).

Registry URL: https://seqco.de/i:57150

**Description of *Bradylimnobacter lacuum* sp. nov.**

*Bradylimnobacter lacuum* (la'cu.um. L. gen. pl. n. *lacuum*, of lakes, referring to the isolation of strains from different lakes).

The type strain is IMCC49206, obtained from Lake Daechung, Republic of Korea, by dilution-to-extinction cultivation in liquid low-nutrient freshwater medium supplemented with bovine catalase. Growth of the type strain occurs at 18 °C, the temperature used for isolation; other cultural properties have not been determined. Another cultured isolate assigned to the same species is IMCC47816, obtained from Lake Paldang. The genome encodes complete pathways for central carbon metabolism and components of aerobic respiration, as well as a proton-pumping rhodopsin, whereas predicted biosynthetic pathways for quinones and heme are incomplete. The genome shares approximately 91% ANI with that of the type strain of *B. soyangensis*, and the DNA G+C content (approximately 33%) is lower than that of *B. exiguus* (approximately 40%). The draft genome assembly of IMCC49206 comprises 2,587,072 bp, with a DNA G+C content of 33.06% and 2,056 predicted genes. The species corresponds to s__UBA961 sp023259575 in GTDB (R226). The nomenclatural type is the genome sequence of strain IMCC49206 (ENA WGS accession number CFWNOQ01).

Registry URL: https://seqco.de/i:57151

**Description of *Phreatorhabdaceae* fam. nov.**

*Phreatorhabdaceae* (Phre.a.to.rhab.da'ce.ae. N.L. fem. n. *Phreatorhabdus*, type genus of the family; L. fem. pl. n. suff. *-aceae*, ending to denote a family; N.L. fem. pl. n. *Phreatorhabdaceae*, the *Phreatorhabdus* family).

The family was previously known only from metagenome-assembled genomes and is represented in GTDB (R226) as the family f__NICIL-2, named after genome NICIL-2. These genomes originate mainly from hot springs and wastewater. Cultivated representatives, obtained from groundwater, are available exclusively as dilution-to-extinction cultures. In GTDB (R226), the family comprises four genera, and *Phreatorhabdus* represents a genus not previously captured in GTDB. The description of the family is otherwise the same as that of the genus *Phreatorhabdus*. The type genus is *Phreatorhabdus*.

Registry URL: https://seqco.de/i:57152

**Description of *Phreatorhabdus* gen. nov.**

*Phreatorhabdus* (Phre.a.to.rhab'dus. Gr. neut. n. *phrear*, a well; Gr. fem. n. *rhabdos*, a rod; N.L. fem. n. *Phreatorhabdus*, a rod from groundwater).

Cells are rods with flagella. Does not form colonies on agar surface. Cultivated representatives have been obtained from groundwater in the Republic of Korea by dilution-to-extinction cultivation and grow in liquid low-nutrient freshwater medium supplemented with bovine catalase. Genome sizes are approximately 4.1 Mbp, with DNA G+C contents of approximately 45.6%. The genus represents an unnamed genus within f__NICIL-2 in GTDB (R226). The type species is *Phreatorhabdus flagellata*.

Registry URL: https://seqco.de/i:57153

**Description of *Phreatorhabdus flagellata* sp. nov.**

*Phreatorhabdus flagellata* **(**fla.gel.la'ta. L. fem. part. adj. *flagellata*, flagellated).

The type strain is IMCC37485, obtained from groundwater of the Yanggu aquifer, Republic of Korea, by dilution-to-extinction cultivation in liquid low-nutrient freshwater medium supplemented with bovine catalase. Cells of the type strain are rods, approximately 0.2 × 2.0 µm, with flagella, as observed by electron microscopy. Growth occurs at 4–37 °C (optimum 25–30 °C), with the highest specific growth rate (approximately 0.9 $day^{-1}$) at 25 °C and the maximum cell density (approximately $4.4 \times 10^7$ cells $mL^{-1}$) at 20 °C. At 37 °C, growth occurs only after a lag of approximately 15 days. Additional cultured isolates assigned to the same species are IMCC31542, IMCC37487, IMCC37488, IMCC37489, IMCC37493, IMCC37499, IMCC37500, and IMCC37502, obtained from the same aquifer. Among them, IMCC31542 grows more slowly (approximately 0.6 $day^{-1}$) and shows no detectable growth at 37 °C. The genome encodes complete pathways for central carbon metabolism and components of aerobic respiration, as well as flagellar genes, respiratory nitrate reductase (*narGH*), and nitrous oxide reductase (*nosZDFYL*), whereas predicted biosynthetic pathways for quinones and heme are incomplete. The genome (approximately 4.1 Mbp) is the largest among the cultivated members of the order *Kapaibacteriales*. The draft genome assembly of IMCC37485 comprises 4,058,958 bp, with a DNA G+C content of 45.56% and 3,364 predicted genes. The species belongs to f__NICIL-2 but is not assigned to any genus or species in GTDB (R226). The nomenclatural type is the genome sequence of strain IMCC37485 (ENA WGS accession number CFWNPP01).

Registry URL: https://seqco.de/i:57154

### Supplementary figures

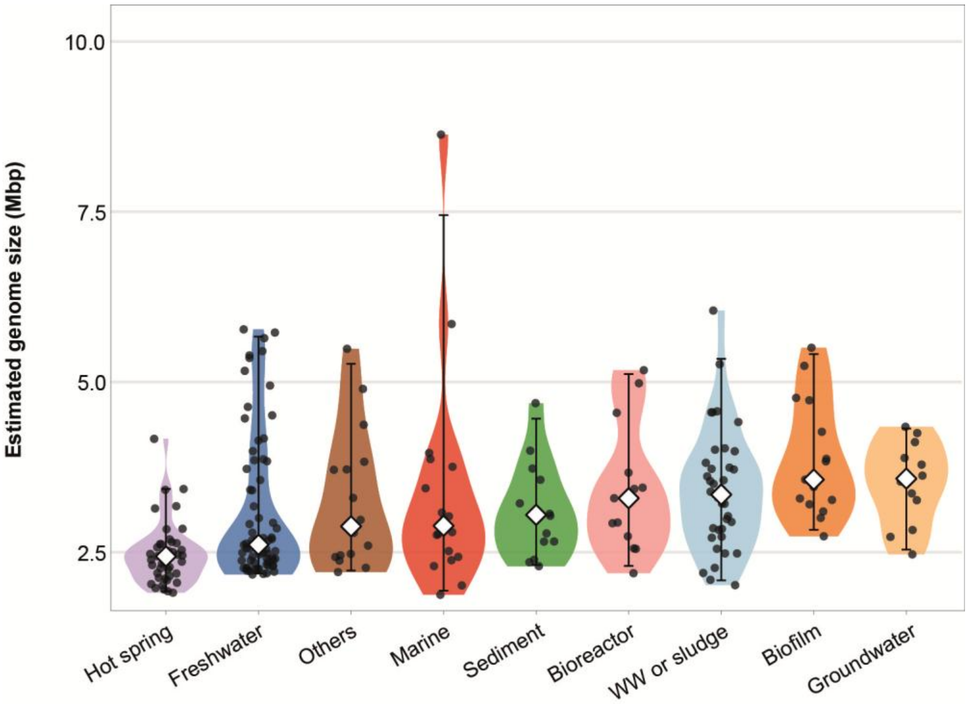


**Fig. S1. Genome size variation of high-quality *Kapaibacteriales* MAGs across habitats.** Estimated genome size was calculated as genome size × 100 / completeness. Points represent individual MAGs (Table S5). In each violin plot, the white diamond indicates the median, and the vertical bar indicates the central 95% of values (2.5th–97.5th percentiles; median_hilow). Habitats are ordered by increasing median genome size. Major habitats represented by fewer than five MAGs are not shown. WW, wastewater

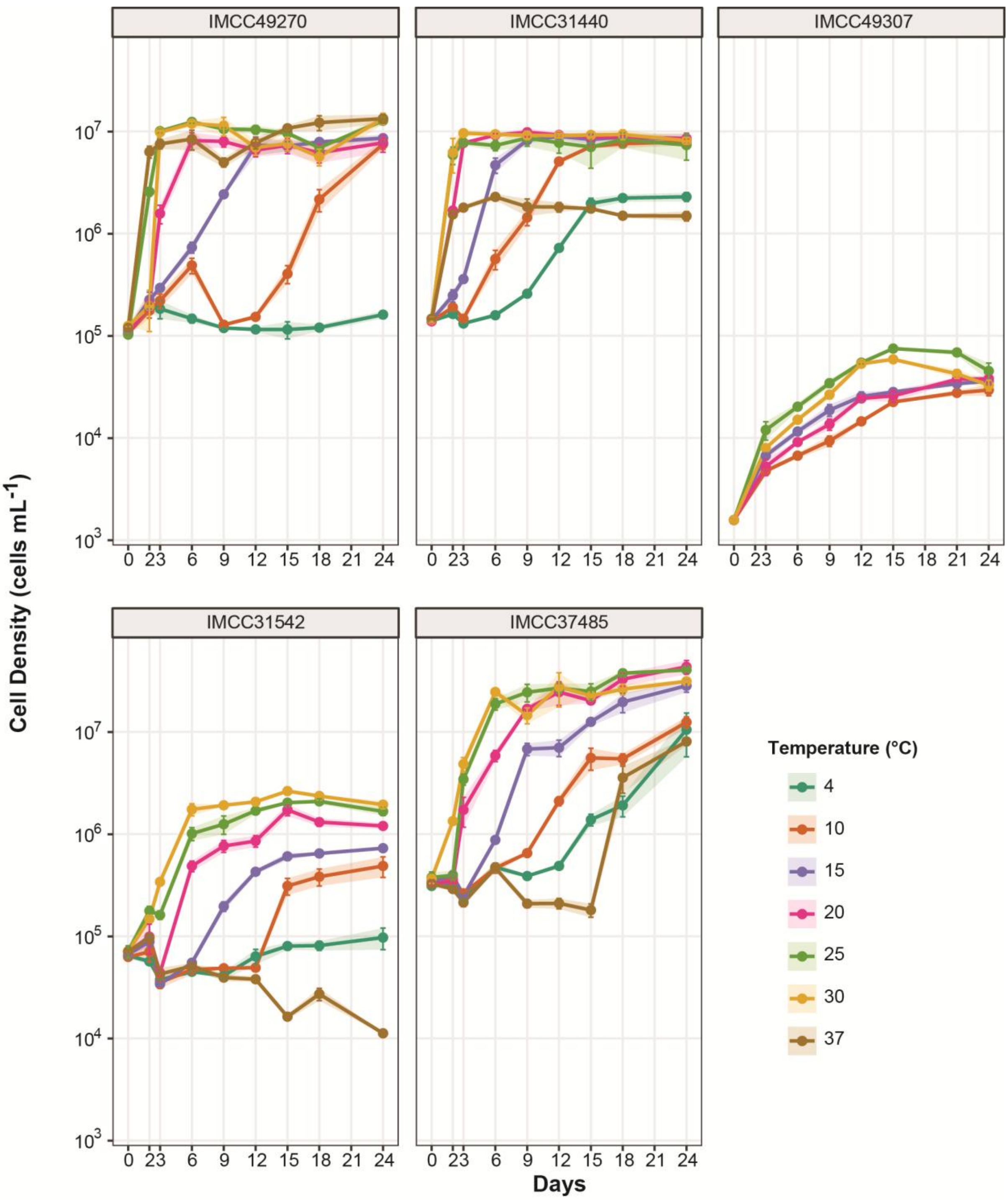


**Fig. S2. Temperature-dependent time-course growth curves of five representative *Kapaibacteriales* isolates.** Time-series growth profiles of five isolates across a range of incubation temperatures (4–37 °C) over 24 days. Cell density was measured periodically at indicated time points. Line colors denote different temperature conditions (4, 10, 15, 20, 25, 30, and 37 °C).

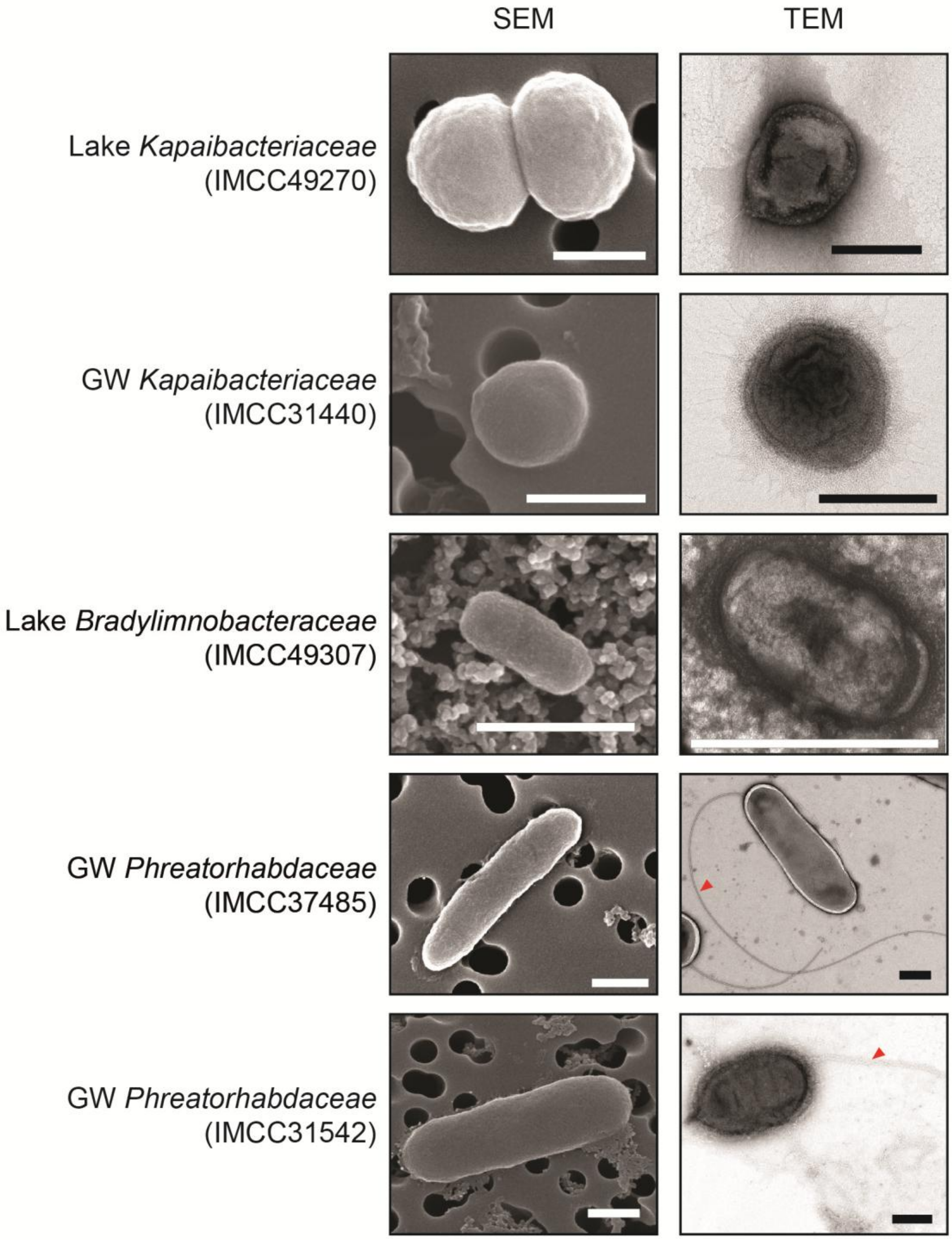


**Fig. S3. Cellular morphology of five representative *Kapaibacteriales* isolates**. Scanning electron microscopy (SEM; left) and transmission electron microscopy (TEM; right) images of five representative isolates. White and black scale bars represent 0.5 μm. Red arrowheads indicate flagella. GW, groundwater

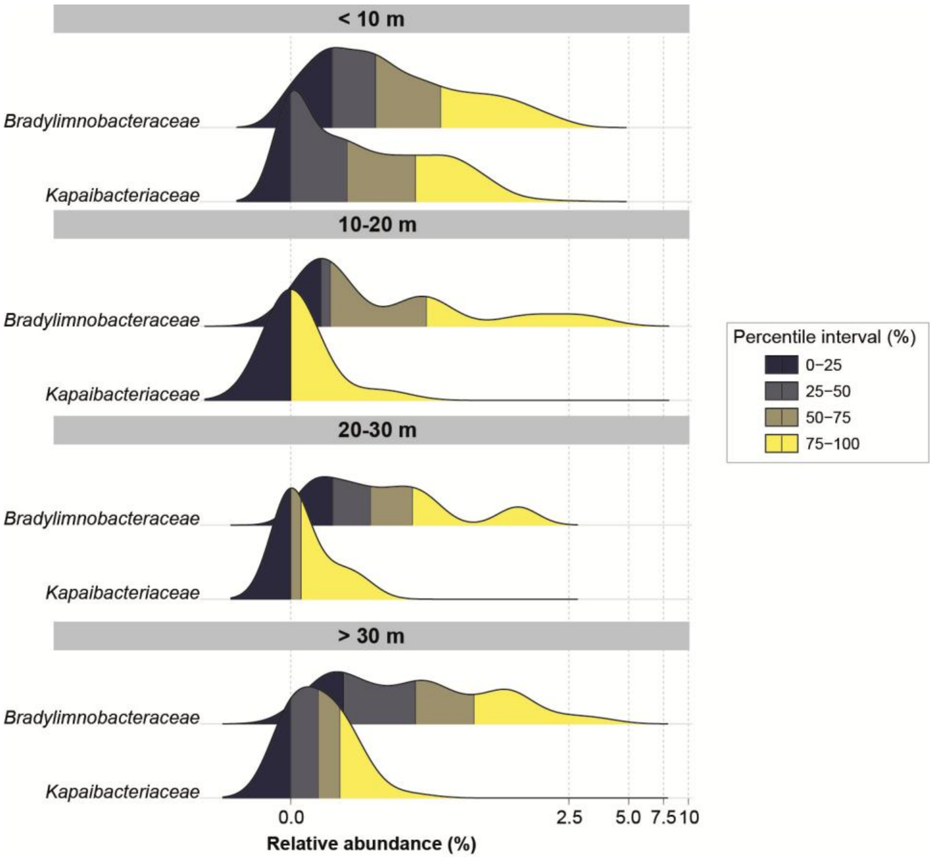


**Fig. S4. Depth-associated distribution of *Bradylimnobacteraceae* and *Kapaibacteriaceae* in lakes.** Ridgeline density plots showing the relative abundance distributions (%) of *Bradylimnobacteraceae* and *Kapaibacteriaceae* in 2,067 metagenomes, grouped into four depth strata (< 10 m, 10–20 m, 20–30 m, and > 30 m). Relative abundances are estimated using SingleM. Shading distinguishes the four percentile intervals of the relative abundance distribution (0–25th, 25th–50th, 50th–75th, and 75th–100th percentiles).

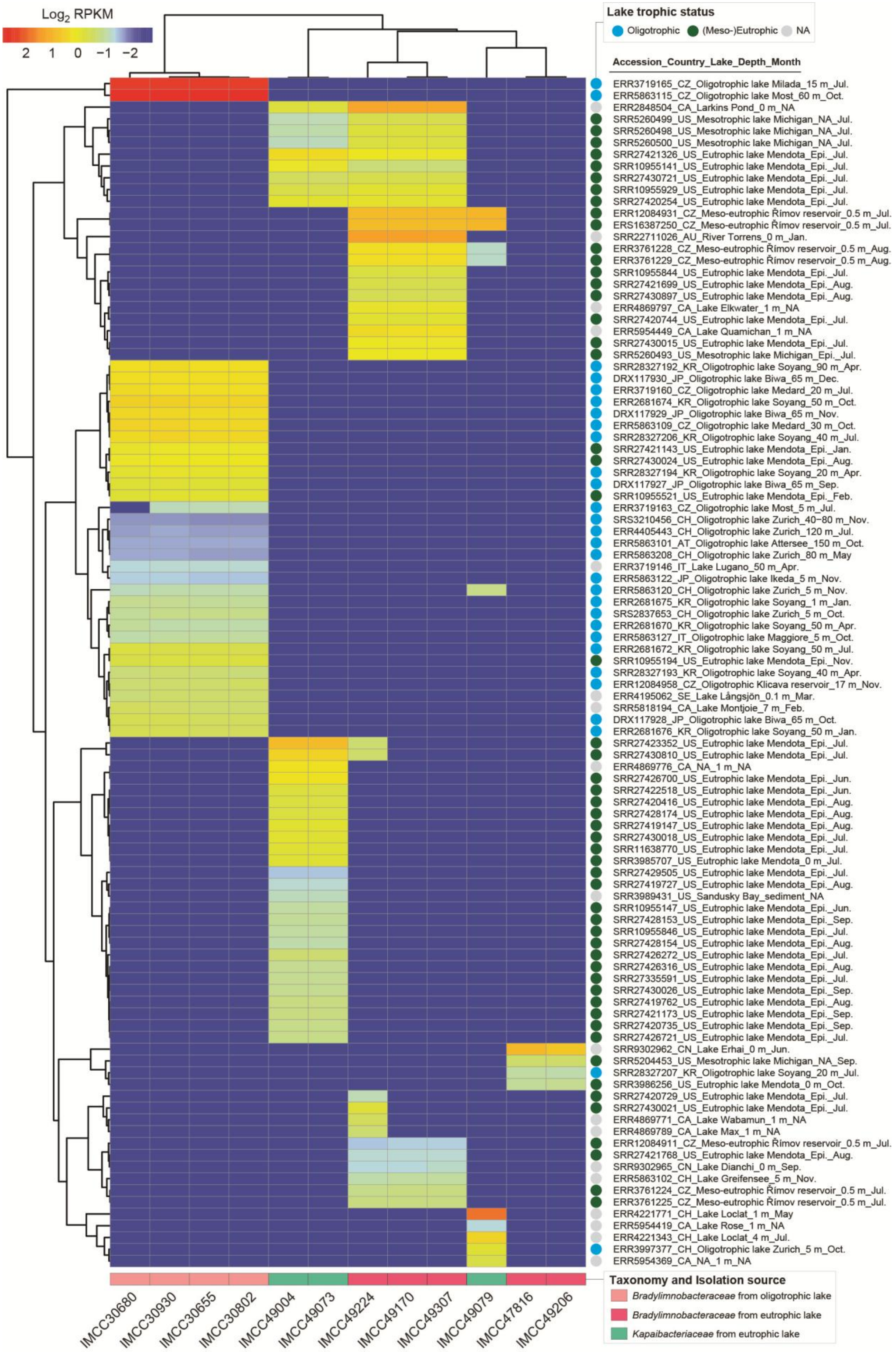

Log2 RPKM
2 1 0 −1 −2
Lake trophic status
Oligotrophic
(Meso-)Eutrophic
NA
Accession_Country_Lake_Depth_Month
ERR3719165_CZ_Oligotrophic lake Milada_15 m_Jul.
ERR5863115_CZ_Oligotrophic lake Most_60 m_Oct.
ERR2848504_CA_Larkins Pond_0 m_NA
SRR5260499_US_Mesotrophic lake Michigan_NA_Jul.
SRR5260498_US_Mesotrophic lake Michigan_NA_Jul.
SRR5260500_US_Mesotrophic lake Michigan_NA_Jul.
SRR27421326_US_Eutrophic lake Mendota_Epi._Jul.
SRR10955141_US_Eutrophic lake Mendota_Epi._Jul.
SRR27430721_US_Eutrophic lake Mendota_Epi._Jul.
SRR10955929_US_Eutrophic lake Mendota_Epi._Jul.
SRR27420254_US_Eutrophic lake Mendota_Epi._Jul.
ERR12084931_CZ_Meso-eutrophic Římov reservoir_0.5 m_Jul.
ERS16387250_CZ_Meso-eutrophic Římov reservoir_0.5 m_Jul.
SRR22711026_AU_River Torrens_0 m_Jan.
ERR3761228_CZ_Meso-eutrophic Římov reservoir_0.5 m_Aug.
ERR3761229_CZ_Meso-eutrophic Římov reservoir_0.5 m_Aug.
SRR10955844_US_Eutrophic lake Mendota_Epi._Jul.
SRR27421699_US_Eutrophic lake Mendota_Epi._Aug.
SRR27430897_US_Eutrophic lake Mendota_Epi._Aug.
ERR4869797_CA_Lake Elkwater_1 m_NA
SRR27420744_US_Eutrophic lake Mendota_Epi._Jul.
ERR5954449_CA_Lake Quamichan_1 m_NA
SRR27430015_US_Eutrophic lake Mendota_Epi._Jul.
SRR5260493_US_Mesotrophic lake Michigan_Epi._Jul.
SRR28327192_KR_Oligotrophic lake Soyang_90 m_Apr.
DRX117930_JP_Oligotrophic lake Biwa_65 m_Dec.
ERR3719160_CZ_Oligotrophic lake Medard_20 m_Jul.
ERR2681674_KR_Oligotrophic lake Soyang_50 m_Oct.
DRX117929_JP_Oligotrophic lake Biwa_65 m_Nov.
ERR5863109_CZ_Oligotrophic lake Medard_30 m_Oct.
SRR28327206_KR_Oligotrophic lake Soyang_40 m_Jul.
SRR27421143_US_Eutrophic lake Mendota_Epi._Jan.
SRR27430024_US_Eutrophic lake Mendota_Epi._Aug.
SRR28327194_KR_Oligotrophic lake Soyang_20 m_Apr.
DRX117927_JP_Oligotrophic lake Biwa_65 m_Sep.
SRR10955521_US_Eutrophic lake Mendota_Epi._Feb.
ERR3719163_CZ_Oligotrophic lake Most_5 m_Jul.
SRS3210456_CH_Oligotrophic lake Zurich_40−80 m_Nov.
ERR4405443_CH_Oligotrophic lake Zurich_120 m_Jul.
ERR5863101_AT_Oligotrophic lake Attersee_150 m_Oct.
ERR5863208_CH_Oligotrophic lake Zurich_80 m_May
ERR3719146_IT_Lake Lugano_50 m_Apr.
ERR5863122_JP_Oligotrophic lake Ikeda_5 m_Nov.
ERR5863120_CH_Oligotrophic lake Zurich_5 m_Nov.
ERR2681675_KR_Oligotrophic lake Soyang_1 m_Jan.
SRS2837653_CH_Oligotrophic lake Zurich_5 m_Oct.
ERR2681670_KR_Oligotrophic lake Soyang_50 m_Apr.
ERR5863127_IT_Oligotrophic lake Maggiore_5 m_Oct.
ERR2681672_KR_Oligotrophic lake Soyang_50 m_Jul.
SRR10955194_US_Eutrophic lake Mendota_Epi._Nov.
SRR28327193_KR_Oligotrophic lake Soyang_40 m_Apr.
ERR12084958_CZ_Oligotrophic Klicava reservoir_17 m_Nov.
ERR4195062_SE_Lake Långsjön_0.1 m_Mar.
SRR5818194_CA_Lake Montjoie_7 m_Feb.
DRX117928_JP_Oligotrophic lake Biwa_65 m_Oct.
ERR2681676_KR_Oligotrophic lake Soyang_50 m_Jan.
SRR27423352_US_Eutrophic lake Mendota_Epi._Jul.
SRR27430810_US_Eutrophic lake Mendota_Epi._Jul.
ERR4869776_CA_NA_1 m_NA
SRR27426700_US_Eutrophic lake Mendota_Epi._Jun.
SRR27422518_US_Eutrophic lake Mendota_Epi._Jun.
SRR27420416_US_Eutrophic lake Mendota_Epi._Aug.
SRR27428174_US_Eutrophic lake Mendota_Epi._Aug.
SRR27419147_US_Eutrophic lake Mendota_Epi._Aug.
SRR27430018_US_Eutrophic lake Mendota_Epi._Jul.
SRR11638770_US_Eutrophic lake Mendota_Epi._Jul.
SRR3985707_US_Eutrophic lake Mendota_0 m_Jul.
SRR27429505_US_Eutrophic lake Mendota_Epi._Jul.
SRR27419727_US_Eutrophic lake Mendota_Epi._Aug.
SRR3989431_US_Sandusky Bay_sediment_NA
SRR10955147_US_Eutrophic lake Mendota_Epi._Jun.
SRR27428153_US_Eutrophic lake Mendota_Epi._Sep.
SRR10955846_US_Eutrophic lake Mendota_Epi._Jul.
SRR27428154_US_Eutrophic lake Mendota_Epi._Aug.
SRR27426272_US_Eutrophic lake Mendota_Epi._Jul.
SRR27426316_US_Eutrophic lake Mendota_Epi._Aug.
SRR27335591_US_Eutrophic lake Mendota_Epi._Jul.
SRR27430026_US_Eutrophic lake Mendota_Epi._Sep.
SRR27419762_US_Eutrophic lake Mendota_Epi._Aug.
SRR27421173_US_Eutrophic lake Mendota_Epi._Sep.
SRR27420735_US_Eutrophic lake Mendota_Epi._Sep.
SRR27426721_US_Eutrophic lake Mendota_Epi._Jul.
SRR9302962_CN_Lake Erhai_0 m_Jun.
SRR5204453_US_Mesotrophic lake Michigan_NA_Sep.
SRR28327207_KR_Oligotrophic lake Soyang_20 m_Jul.
SRR3986256_US_Eutrophic lake Mendota_0 m_Oct.
SRR27420729_US_Eutrophic lake Mendota_Epi._Jul.
SRR27430021_US_Eutrophic lake Mendota_Epi._Jul.
ERR4869771_CA_Lake Wabamun_1 m_NA
ERR4869789_CA_Lake Max_1 m_NA
ERR12084911_CZ_Meso-eutrophic Římov reservoir_0.5 m_Jul.
SRR27421768_US_Eutrophic lake Mendota_Epi._Aug.
SRR9302965_CN_Lake Dianchi_0 m_Sep.
ERR5863102_CH_Lake Greifensee_5 m_Nov.
ERR3761224_CZ_Meso-eutrophic Římov reservoir_0.5 m_Jul.
ERR3761225_CZ_Meso-eutrophic Římov reservoir_0.5 m_Jul.
ERR4221771_CH_Lake Loclat_1 m_May
ERR5954419_CA_Lake Rose_1 m_NA
ERR4221343_CH_Lake Loclat_4 m_Jul.
ERR3997377_CH_Oligotrophic lake Zurich_5 m_Oct.
ERR5954369_CA_NA_1 m_NA
IMCC30680
IMCC30930
IMCC30655
IMCC30802
IMCC49004
IMCC49073
IMCC49224
IMCC49170
IMCC49307
IMCC49079
IMCC47816
IMCC49206
Taxonomy and Isolation source
Bradylimnobacteraceae from oligotrophic lake
Bradylimnobacteraceae from eutrophic lake
Kapaibacteriaceae from eutrophic lake

**Fig. S5. Metagenomic read recruitment to genomes of lake-derived *Kapaibacteriales* isolates.** Heatmap showing the abundance of isolate genome (columns) in each metagenome (rows). A genome was considered present in a metagenome when recruited reads covered at least 50% of its length (covered fraction $\geq 0.5$). Only genomes detected in at least one metagenome and metagenomes with at least one detected genome are shown. Hierarchical clustering dendrograms reflect similarities in recruitment profiles across genomes and metagenomic samples. NA, not available; Epi., epilimnion.

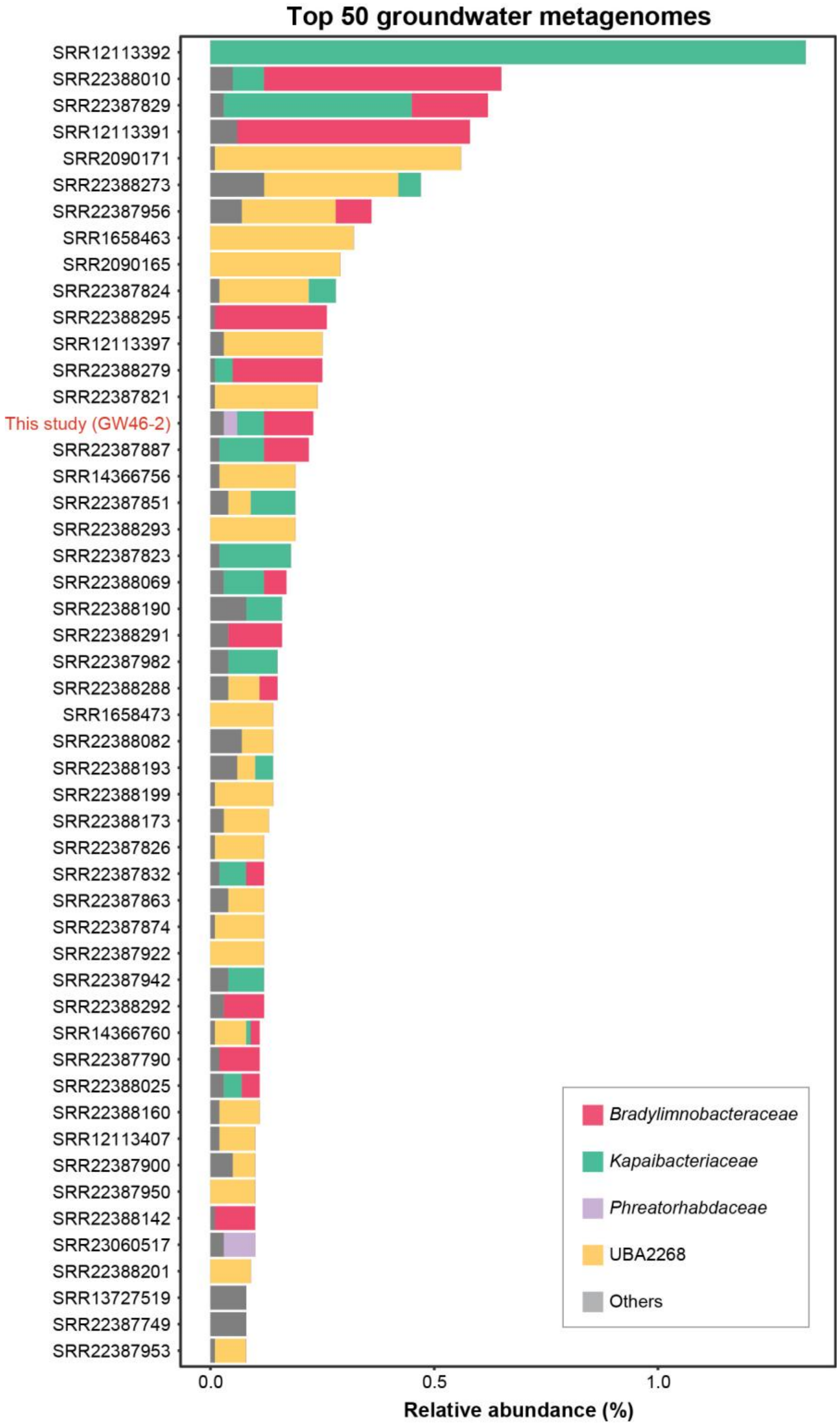


**Fig. S6. Family-level composition of *Kapaibacteriales* in groundwater metagenomes.** Stacked bars show relative abundances (%) estimated using SingleM for the 50 groundwater metagenomes with the highest total *Kapaibacteriales* relative abundance, ordered from highest to lowest. Colors distinguish families, with remaining families grouped as 'Others'. The groundwater metagenome generated in this study (GW46-2) is highlighted in red.

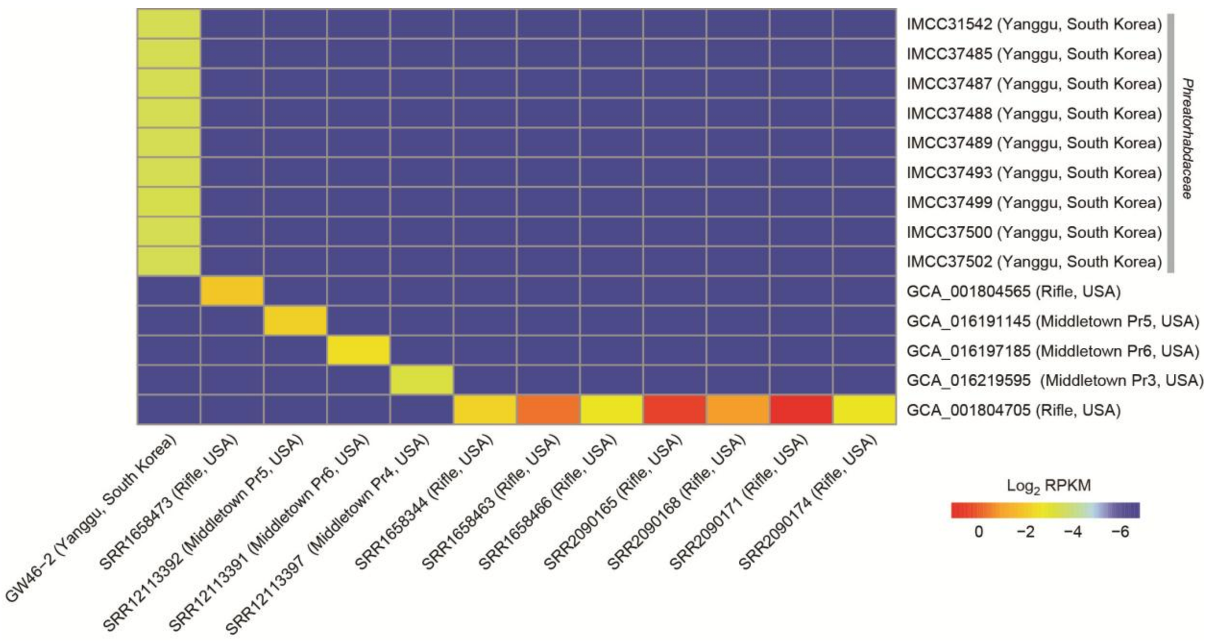


**Fig. S7. Metagenomic read recruitment to *Kapaibacteriales* groundwater isolate genomes and MAGs.** Heatmap showing the abundance of each isolate genome and MAG (rows) in each metagenome (columns). A genome was considered present in a metagenome when recruited reads covered at least 50% of its length (covered fraction ≥ 0.5). Only genomes detected in at least one metagenome and metagenomes with at least one detected genome are shown.

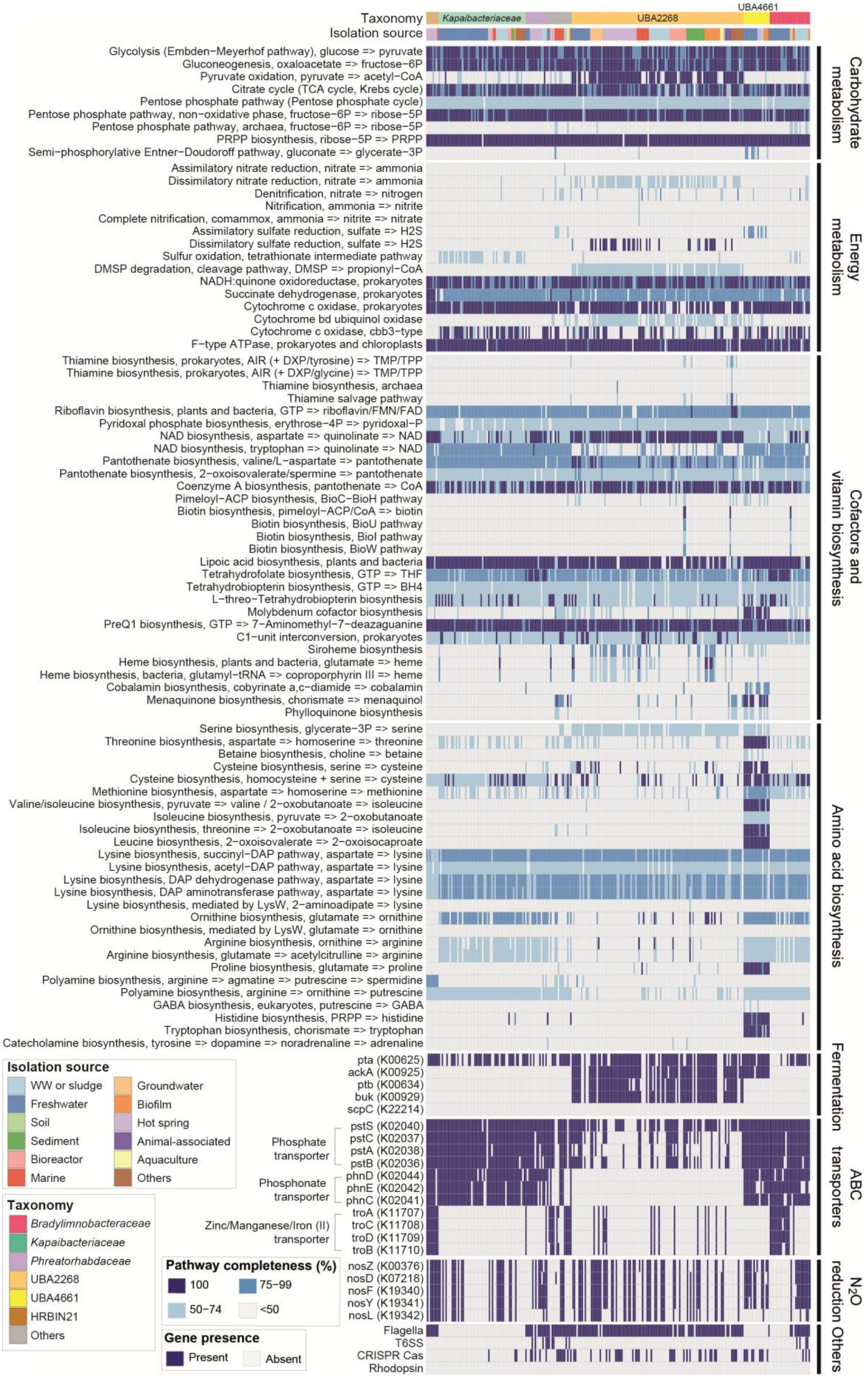

Taxonomy
Isolation source
Kapaibacteriaceae
UBA2268
UBA4661
Glycolysis (Embden−Meyerhof pathway), glucose => pyruvate
Gluconeogenesis, oxaloacetate => fructose−6P
Pyruvate oxidation, pyruvate => acetyl−CoA
Citrate cycle (TCA cycle, Krebs cycle)
Pentose phosphate pathway (Pentose phosphate cycle)
Pentose phosphate pathway, non−oxidative phase, fructose−6P => ribose−5P
Pentose phosphate pathway, archaea, fructose−6P => ribose−5P
PRPP biosynthesis, ribose−5P => PRPP
Semi−phosphorylative Entner−Doudoroff pathway, gluconate => glycerate−3P
Carbohydrate metabolism
Assimilatory nitrate reduction, nitrate => ammonia
Dissimilatory nitrate reduction, nitrate => ammonia
Denitrification, nitrate => nitrogen
Nitrification, ammonia => nitrite
Complete nitrification, comammox, ammonia => nitrite => nitrate
Assimilatory sulfate reduction, sulfate => H2S
Dissimilatory sulfate reduction, sulfate => H2S
Sulfur oxidation, tetrathionate intermediate pathway
DMSP degradation, cleavage pathway, DMSP => propionyl−CoA
NADH:quinone oxidoreductase, prokaryotes
Succinate dehydrogenase, prokaryotes
Cytochrome c oxidase, prokaryotes
Cytochrome bd ubiquinol oxidase
Cytochrome c oxidase, cbb3−type
F−type ATPase, prokaryotes and chloroplasts
Energy metabolism
Thiamine biosynthesis, prokaryotes, AIR (+ DXP/tyrosine) => TMP/TPP
Thiamine biosynthesis, prokaryotes, AIR (+ DXP/glycine) => TMP/TPP
Thiamine biosynthesis, archaea
Thiamine salvage pathway
Riboflavin biosynthesis, plants and bacteria, GTP => riboflavin/FMN/FAD
Pyridoxal phosphate biosynthesis, erythrose−4P => pyridoxal−P
NAD biosynthesis, aspartate => quinolinate => NAD
NAD biosynthesis, tryptophan => quinolinate => NAD
Pantothenate biosynthesis, valine/L−aspartate => pantothenate
Pantothenate biosynthesis, 2−oxoisovalerate/spermine => pantothenate
Coenzyme A biosynthesis, pantothenate => CoA
Pimeloyl−ACP biosynthesis, BioC−BioH pathway
Biotin biosynthesis, pimeloyl−ACP/CoA => biotin
Biotin biosynthesis, BioU pathway
Biotin biosynthesis, BioI pathway
Biotin biosynthesis, BioW pathway
Lipoic acid biosynthesis, plants and bacteria
Tetrahydrofolate biosynthesis, GTP => THF
Tetrahydrobiopterin biosynthesis, GTP => BH4
L−threo−Tetrahydrobiopterin biosynthesis
Molybdenum cofactor biosynthesis
PreQ1 biosynthesis, GTP => 7−Aminomethyl−7−deazaguanine
C1−unit interconversion, prokaryotes
Siroheme biosynthesis
Heme biosynthesis, plants and bacteria, glutamate => heme
Heme biosynthesis, bacteria, glutamyl−tRNA => coproporphyrin III => heme
Cobalamin biosynthesis, cobyrinate a,c−diamide => cobalamin
Menaquinone biosynthesis, chorismate => menaquinol
Phylloquinone biosynthesis
Cofactors and vitamin biosynthesis
Serine biosynthesis, glycerate−3P => serine
Threonine biosynthesis, aspartate => homoserine => threonine
Betaine biosynthesis, choline => betaine
Cysteine biosynthesis, serine => cysteine
Cysteine biosynthesis, homocysteine + serine => cysteine
Methionine biosynthesis, aspartate => homoserine => methionine
Valine/isoleucine biosynthesis, pyruvate => valine / 2−oxobutanoate => isoleucine
Isoleucine biosynthesis, pyruvate => 2−oxobutanoate
Isoleucine biosynthesis, threonine => 2−oxobutanoate => isoleucine
Leucine biosynthesis, 2−oxoisovalerate => 2−oxoisocaproate
Lysine biosynthesis, succinyl−DAP pathway, aspartate => lysine
Lysine biosynthesis, acetyl−DAP pathway, aspartate => lysine
Lysine biosynthesis, DAP dehydrogenase pathway, aspartate => lysine
Lysine biosynthesis, DAP aminotransferase pathway, aspartate => lysine
Lysine biosynthesis, mediated by LysW, 2−aminoadipate => lysine
Ornithine biosynthesis, glutamate => ornithine
Ornithine biosynthesis, mediated by LysW, glutamate => ornithine
Arginine biosynthesis, ornithine => arginine
Arginine biosynthesis, glutamate => acetylcitrulline => arginine
Proline biosynthesis, glutamate => proline
Polyamine biosynthesis, arginine => agmatine => putrescine => spermidine
Polyamine biosynthesis, arginine => ornithine => putrescine
GABA biosynthesis, eukaryotes, putrescine => GABA
Histidine biosynthesis, PRPP => histidine
Tryptophan biosynthesis, chorismate => tryptophan
Catecholamine biosynthesis, tyrosine => dopamine => noradrenaline => adrenaline
Amino acid biosynthesis
pta (K00625)
ackA (K00925)
ptb (K00634)
buk (K00929)
scpC (K22214)
Fermentation
Phosphate transporter
pstS (K02040)
pstC (K02037)
pstA (K02038)
pstB (K02036)
Phosphonate transporter
phnD (K02044)
phnE (K02042)
phnC (K02041)
Zinc/Manganese/Iron (II) transporter
troA (K11707)
troC (K11708)
troD (K11709)
troB (K11710)
ABC transporters
nosZ (K00376)
nosD (K07218)
nosF (K19340)
nosY (K19341)
nosL (K19342)
N2O reduction
Flagella
T6SS
CRISPR Cas
Rhodopsin
Others
Isolation source
WW or sludge
Freshwater
Soil
Sediment
Bioreactor
Marine
Groundwater
Biofilm
Hot spring
Animal-associated
Aquaculture
Others
Taxonomy
Bradylimnobacteraceae
Kapaibacteriaceae
Phreatorhabdaceae
UBA2268
UBA4661
HRBIN21
Others
Pathway completeness (%)
100
75−99
50−74
<50
Gene presence
Present
Absent

**Fig. S8. Predicted metabolic potential of *Kapaibacteriales* MAGs from diverse habitats.** Columns represent individual MAGs, and rows represent KEGG modules or additional functional genes and systems. Pathway completeness is grouped into four categories (100%, 75–99%, 50–74%, and <50%). Additional genes and systems are shown as present or absent according to the genomic annotations. Top annotations indicate family-level taxonomy and source habitat. Functional categories are indicated on the right. T6SS, type VI secretion system.

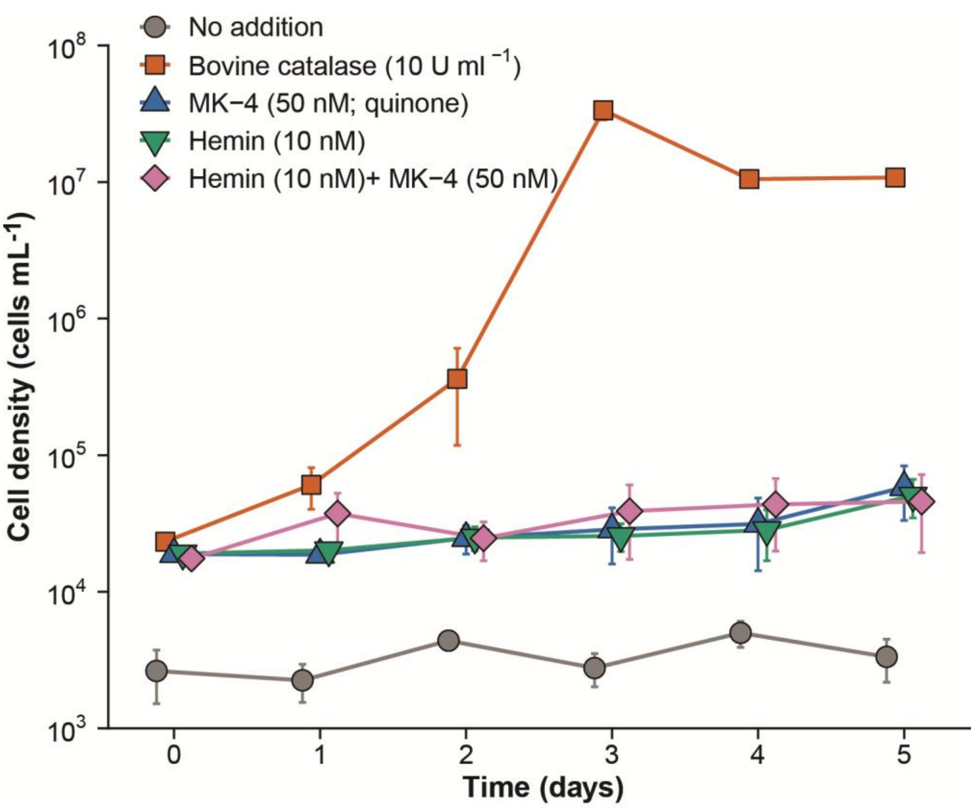


**Fig. S9. Growth of IMCC31440 under catalase, hemin, and quinone supplementation.** Strain IMCC31440 was grown in LNFM with no supplementation, or with catalase (10 U $mL^{-1}$), menaquinone-4 (MK-4, 50 nM), hemin (10 nM), or hemin plus MK-4 (10 nM and 50 nM, respectively). Cultures were incubated at 25 °C and cell densities were determined by flow cytometry. Symbols represent means of three biological replicates and error bars show the standard deviations.

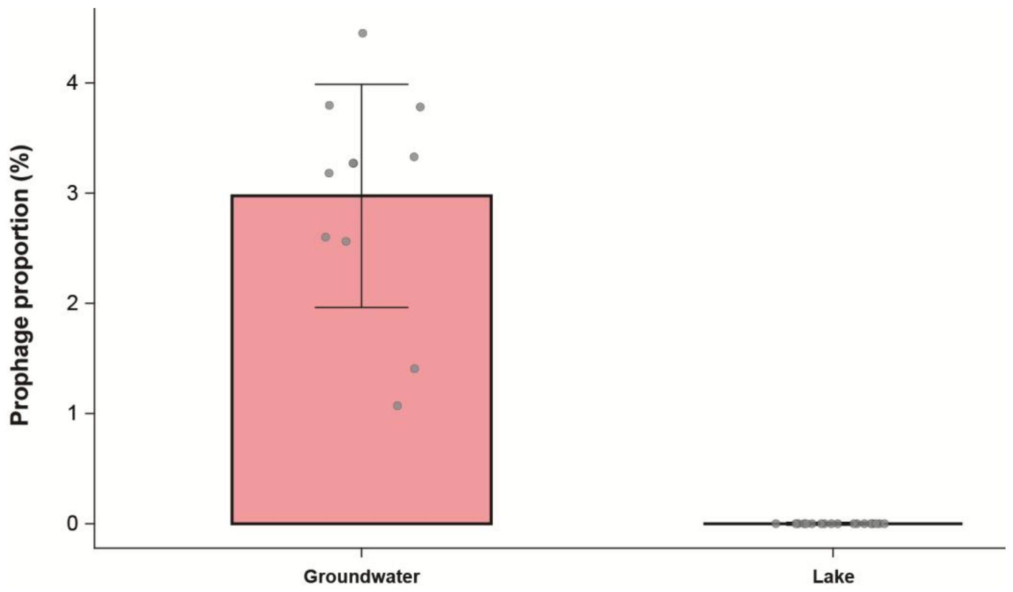


**Fig. S10. Prophage proportions in *Kapaibacteriales* isolate genomes (*n* = 29) from groundwater and lakes.** Bars show the mean prophage proportion (%) for each habitat group, and error bars indicate standard deviations. Grey points represent individual isolate genomes.

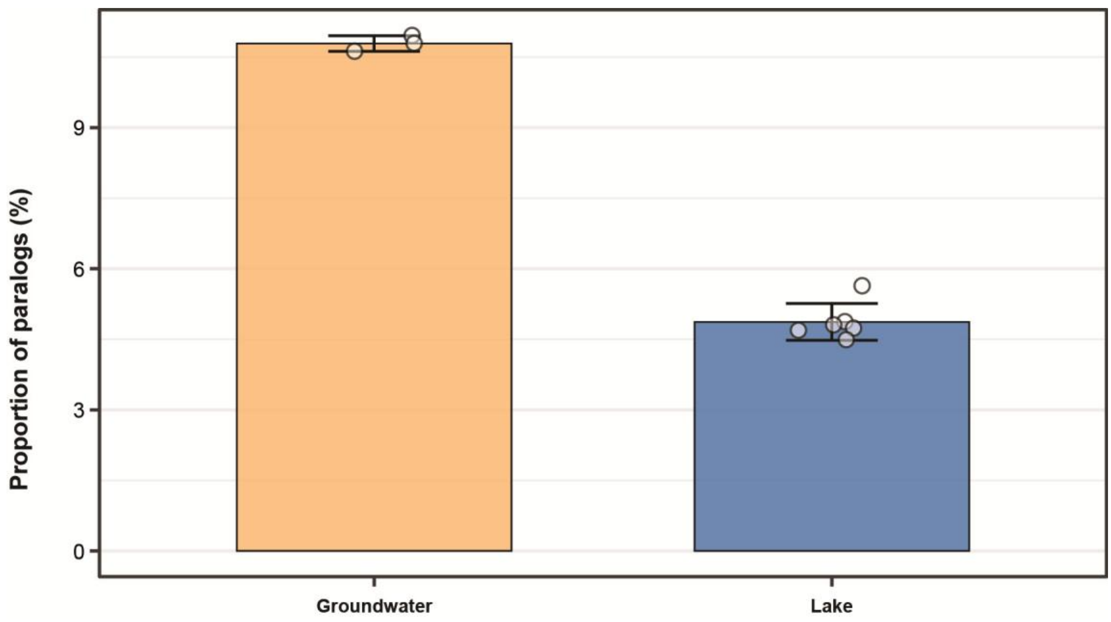


**Fig. S11. Paralog proportions in *Kapaibacteriales* species from groundwater and lakes.** Analyses used one representative genome per species, comprising three groundwater and six lake species. Bars show the mean percentage of genes classified as paralogs in each habitat group, and error bars indicate standard deviations. Open circles represent individual species.

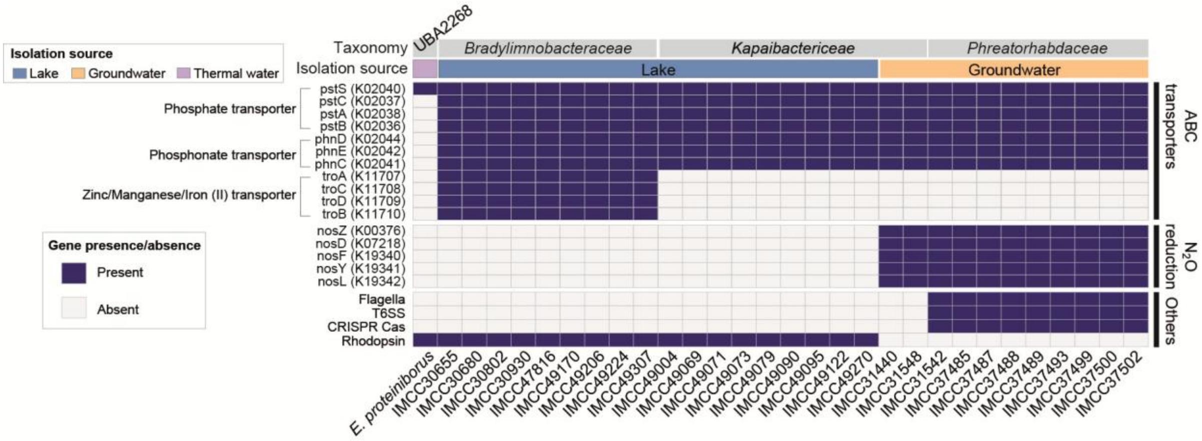


**Fig. S12. Distribution of selected functional genes and systems across *Kapaibacteriales* isolates.** Columns represent isolate genomes and rows represent genes or functional systems. Top annotations indicate family-level taxonomy and isolation source. Functional categories are indicated on the right. T6SS, type VI secretion system.
